# Early life γδ T cell activation enforces intestinal barrier integrity during intergenerational *C. difficile* colonization

**DOI:** 10.64898/2026.04.28.721483

**Authors:** Bavanitha Thurairajah, Susan Westfall, Lisa Chiggiato, Liam Keogh, Anshul Sinha, Kaitlin A. Olsen, Christina Gavino, Yohannes Tafesse, Emmanuelle Moens, Patrick Lypaczewski, Ghislaine Fontes, B. Jesse Shapiro, Benoit Cousineau, Samantha Gruenheid, David Vermijlen, Bastien Castagner, Antoine-Emmanuel Saliba, Irah L. King

## Abstract

Early postnatal life is a highly dynamic period in which the intestine is inundated with billions of microbes that not only battle for colonization, but also strongly influence immune system function and long-term health. This brief period of development provides a window of opportunity to imprint resilience against disease, yet the intercellular dialogue that dictates tissue protection are only beginning to be discovered. By performing a kinetic analysis of immune cell activation in the colon of mice from early life to adulthood, we identified transient activation of a microbiota-dependent type 3 immune response during the weaning period. This response was characterized by a selective increase in IL-17 production by fetal-derived γδ T cells that occurred in an IL-1 receptor-dependent, cell-intrinsic manner. The subsequent differentiation of IL-10 producing Rorγt+ T regulatory cells extinguished IL-17 production by γδ T cells and prevented immunopathology. Microbial gain-of-function approaches determined that colonic γδ T cell activation occurred in response to passive acquisition of *C. difficile* at birth. In turn, IL-17 limited *C. difficile* growth and intestinal barrier breach by the commensal microbiota. Collectively, our results reveal how an agile immune cell network meets the demands of a maturing microbiota to support gut health during early life.

**Highlights:**

- Colonic γδ17 T cells are activated during early life in a microbiota-specific manner
- Fetal-derived γδ17 T cells require IL-1 signals for activation
- Passive acquisition of *C.difficile* induces γδ17 T cell activation
- IL-17 limits *C. difficile* growth and systemic dissemination of commensals

## INTRODUCTION

As proposed by Herzenberg and Herzenberg, the layered immune hypothesis suggests that lymphocyte repertoires develop by distinct waves of lineage commitment over the life span^1^. During early life, these repertoires are enriched in innate-like lymphocyte populations that are thought to provide non-specific immune defense prior to the development of antigen-specific lymphocytes and regulatory cell types. While this paradigm was developed from rodent studies, they have recently been validated in human studies^2^. Of the innate-like lymphocyte subsets, γδ T cells have the earliest ontogeny, predominantly emerging from the thymus between late gestation and shortly after birth^3^. During this wave of development, reception of differential T cell receptor signaling imprints γδ T cells with distinct effector programs^4^. As such, γδ T cells are categorized by their Vγ chain usage^5^ and effector functions^6^. In mice, monoclonal Vγ5+ and Vγ7+ γδ T cell subsets residing within the most superficial layers of the skin and intestine, respectively, maintain an intricate dialogue with epithelial cells by responding to self and producing growth factors that regulate barrier homeostasis, anti-cancer immunity and wound healing^7^. In contrast, Vγ4+ and Vγ6+ γδ T cells express the transcription factor Rorγt, endowing them with the ability to produce type 3 cytokines such as IL-17 and IL-22, whereas Vγ1+ thymocytes express Tbet and are poised to produce IFNγ upon activation^8^.

Within the γδ T cell lineage, IL-17 producing γδ T cells (hereafter referred to as γδ17 T cells) are unique in that they are prevalent in all barrier tissues and been demonstrated to perform pleiotropic roles in host defense and tissue physiology^9^. For example, γδ17 T cells mediate anti-fungal^10^ and anti-bacterial^11^ immunity as well as promote wound healing^12^, barrier integrity^13^ and metabolic processes such as lipolysis^14^ and epithelial nutrient sensing^15^. However, protracted cytokine production by γδ17 T cells, particularly IL-17, has been implicated in diverse disease states including psoriasiform inflammation^16^, liver disease^17^, fibrosis^18,19^, autoimmunity^20^ and colorectal cancer^21^. Thus, these cells must be tightly regulated to support maximize their protective properties while avoiding tissue pathology.

The diverse functions of γδ17 T cells reflect the multiple stimuli that drive their activation. Indeed, γδ T cells are not MHC-restricted but are activated in an innate-like manner by various environmental stimuli including pathogen-derived products, danger-associated molecular patterns and cytokines^22^. Given their enrichment in barrier sites, it stands to reason that the microbiota has important tissue-specific influences on γδ17 T cell responses^23^. Consistently, γδ17 T cells have been shown to be regulated by commensal microorganisms at the cutaneous^24^, oral^25^, lung^26^, intestinal^27,28^, genital^29^ and ocular^30^ barriers. While these studies were carried out in adult tissues, recent longitudinal studies of human cohorts suggest that γδ T cells expand soon after birth^31,32^ and that this change is associated with an increase in microbiota-driven IL-17 production^32^. Given these results, we hypothesized that γδ17 T cells in the microbial-rich colon are particularly well-positioned to respond to intestinal microbiota changes that occur in early life and have an important function in supporting gut health during this dynamic period of life.

## RESULTS

### Early life activation of colonic γδ17 T cells

Cells poised to produce type 3 cytokines IL-17 and IL-22 dominate the large intestinal lamina propria (LP) under adult, steady-state conditions^33^. However, the repertoire and responsiveness of these cells in the colon during early life is unclear. To address this knowledge gap, we quantified IL-17 and IL-22 production by flow cytometry analysis of CD45+ hematopoietic cells isolated from the colonic LP at various ages of mice. In the absence of in vitro stimulation, we detected a selective increase in IL-17+ cells at 3 weeks of age, whereas we detected no change in IL-22+ cells over time (Figure 1A-C). Notably, the transient change in IL-17 was largely due to a selective increase in production by Rorγt+ γδ T cells in all regions of the colon (Figure 1D-G; Figure S1A). Rorγt+ γδ T cells also increased IL-22 production at weaning, however, Rorγt+ ILC3s were the dominant and stable source of colonic IL-22 at all ages examined (Figure S1B). Dynamic expression of IL-17 by γδ T cells was not due to cell-intrinsic regulatory mechanisms as in vitro stimulation of colonic LP cells with IL-1β and IL-23 showed that Rorγt+ γδ T cells were competent to produce IL-17 and IL-22 at all ages examined (Figure 1H, Figure S1C). Furthermore, colonic IL-17 production was not compensated by other cells in the absence of γδ T cells, whereas IL-22 production was largely unaffected (Figure 1I). IL-17 production by γδ T cells was specific to the colon and undetectable in other barrier tissues such as the small intestine, skin and lungs (Figure 1J). Whole tissue lysates from the colon corroborated our flow cytometry results, indicating that IL-17, but not the type 1 and type 2 cytokines, IFNγ or IL-13, differed at 3 and 8 weeks of age (Figure S1D). A previous study described a transient increase in IFNγ and TNFα in the ileum of weanling mice that they termed the weaning reaction^34^. Consistent with that work, we consistently detected a population of IFNγ-producing CD4+ T cells in the ileum, but not colon, of weanling mice that subsided with age (Figure S1E).

**Figure 1.**
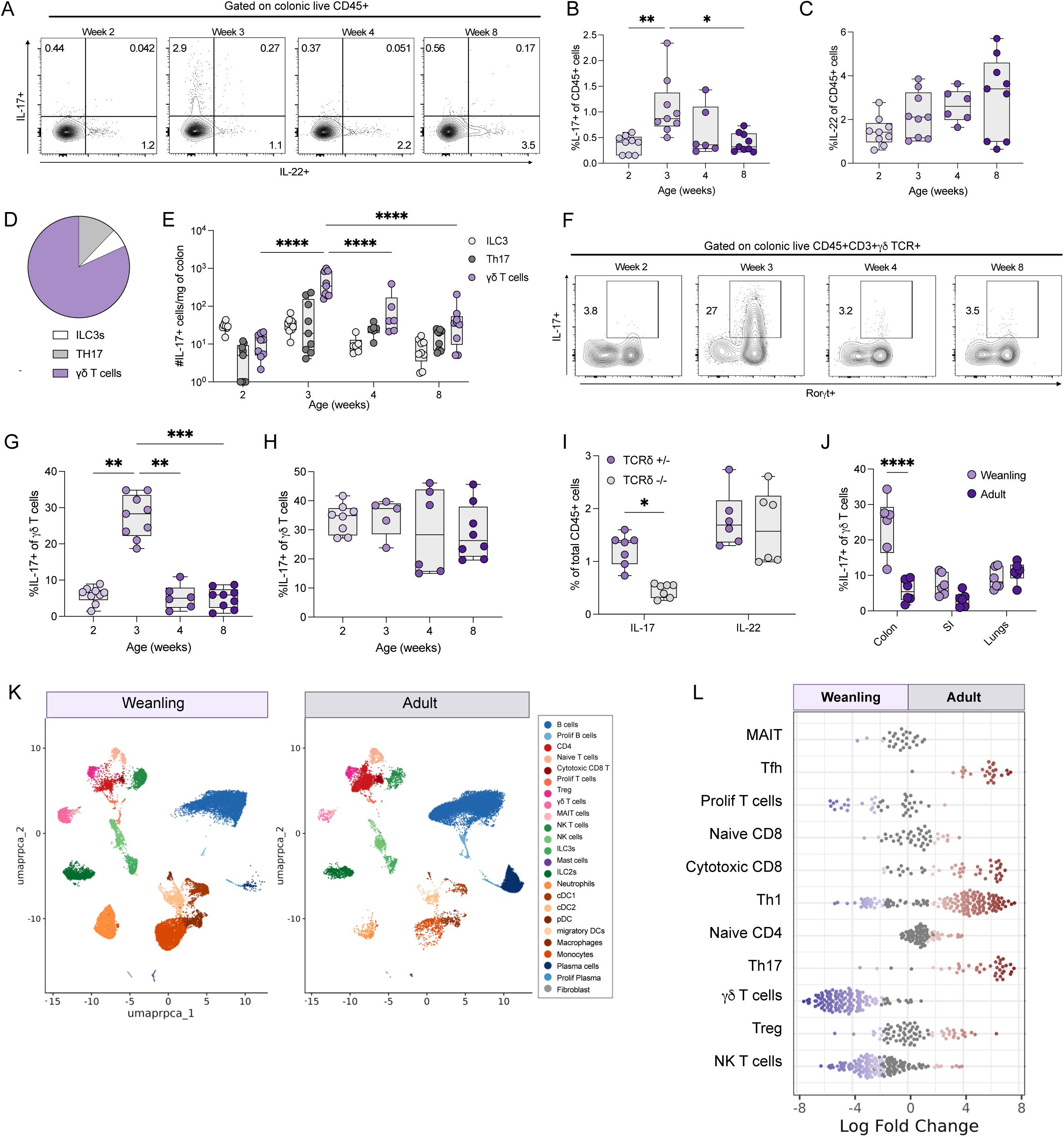
Colonic. γδ**17 T cell activation occurs in early life**. **(A-C)** Representative plots (A) and percent of IL-17+ (B) and IL-22+ (C) of total CD45 cells at indicated ages. **(D-E)** Proportion (D) and number (E) of IL-17+ γδ T cells, TH17 and ILC3s in the colonic lamina propria at indicated times. **(F-G)** Representative plot (F) and percent (G) of IL-17+ of LP γδ T cells of colons of mice at indicated ages. **(H)** Percent of IL-17+ of LP γδ T cells of colons of mice at indicated ages following 4-hour stimulation with 10ng/ml IL-1β and 10ng/ml IL-23. **(I)** Percent of IL-17 and IL-22 of CD45+ cells in colonic LP of δTCR+/-and δTCR-/-mice. **(J)** Percent of IL-17+ γδ T cells in colon, small intestine, lungs and skin in weanling and adult mice. **(K, L)** Single cell UMAP of total CD45+ cells (K) and differential abundance using miloR analysis of total *Cd3*-expressing population of the lamina propria of weanling and adult mice. (B, C, E, and G-J) Each dot represents cells from an individual mouse. Data are represented as mean with SD. Graphs are pooled data from at least two independent experiments with at least n = 3 in each experiment. (A and F) Contour plots are representative at least three independent experiments. Numbers indicate frequency of cells within each quadrant or box. (E, I, J) Data were analyzed using 2 way ANOVA test or (B, C, G, H) Kruskal-Wallis multiple comparison test (∗p < 0.05, ∗∗p < 0.01, ∗∗∗p < 0.001, ∗∗∗∗p <0.0001).

To complement our flow cytometry data, we performed single cell RNA sequencing on CD45+ cells from the colons of 3-and 8-week-old mice (Figure 1K). Barcoded Totalseq anti-TCRδ antibodies were included to ensure that we could distinguish γδ T cells from other T cells lineages (Figure S1F). We found that while LP γδ T cells can be readily identified from other immune populations (based on *Tcrd* transcripts), barcode-labeled antibodies were beneficial in differentiating these cells other innate-like T cells, such as mucosal-associated invariant (MAIT) cells. MiloR analysis indicated numerous differences were observed between these age groups, including that *Tcrd*-expressing cells represented a greater proportion of *Cd3*+ T cells at 3 weeks of age compared to adults (Figure 1L). In adults, the proportions seemed to switch, such that αβ T cells were more prominent, including an increase in *Rorc*-expressing CD4+ Th17 cells (Figure 1L). Colonic γδ T cells in weanling mice expressed genes related to pathways associated with cell-cell adhesion such as *Abi3bp, Tln1, Adam12* and *Itgav* and proliferation such as *Actb*, *Plac8*, *Rbl2*, *Gimap5* and *Pfn1*. *Cd163l1* expression, that codes for Scart-1 and previously shown to be expressed by cutaneous Vγ6+ γδ T cells^35^, was also enriched at weaning (Figure S1G, H). Interestingly, γδ T cells from the adult colon express genes downstream of TCR signaling such as *Socs1, Ctla2a*, and *Lck*. Collectively, these data indicate that colonic γδ17 T cells are selectively activated and express a gene expression signature unique to the weaning period. Given that only IL-17, not IL-22, significantly changed over time, we focused our attention on the mechanisms underlying transient IL-17 production of colonic γδ T cells during early life.

### Cytokine-producing γδ17 T cells are fetal-derived

Previous studies have shown that IL-17 producing γδ T cells develop primarily during the perinatal period between E15.5 and birth^3^. These cells seed peripheral tissues in early-life and are suggested to be maintained within these tissues long-term^36^. To examine this idea further, we assessed Vγ usage by the IL-17-producing γδ T cells in the lamina propria via flow cytometry. Consistent with our scRNAseq results, the majority of colonic LP γδ T cells expressed the Vγ6 subunit at all ages examined, with a minority expressing either Vγ4 or Vγ1 (Figure 2A). Vγ6+ γδ T cells were also the dominant population of IL-17 producing cells at all ages examined, even though Vγ4+ γδ17 T cells also increased IL-17 production at weaning age (Figure 2B, C). Single cell sequencing confirmed that Vγ6+ γδ T cells made up a large proportion of neonatal thymocytes and highly expressed *Il17a* transcripts (Figure 2D-F). To determine whether fetal-derived Vγ6+ γδ T cells can persist in the colon and give rise to γδ17 T cells in weanling mice, we developed an adoptive transfer approach in which thymocytes isolated from 1 day-old pups are transferred into 1 day-old γδ T cell-deficient recipient animals via the temporal vein (Figure 2G). Compared to mice that did not receive 1 day-old thymocytes, we were able to readily detect γδ T cells in the colonic LP, not the IEL fraction, of weanling mice (Figure 2H, I). The donor γδ T cells isolated from weanling recipients were predominantly Vγ6+ (Figure 2J, K). Notably, donor γδ T cells isolated from weanling mice readily produced IL-17, whereas donor cells from adult mice did not (Figure 2L, M). Together, our data indicate that colonic γδ T cells are largely composed of long-lived, fetal-derived Vγ6+ cells that produce IL-17 in an age-dependent manner.

**Figure 2.**
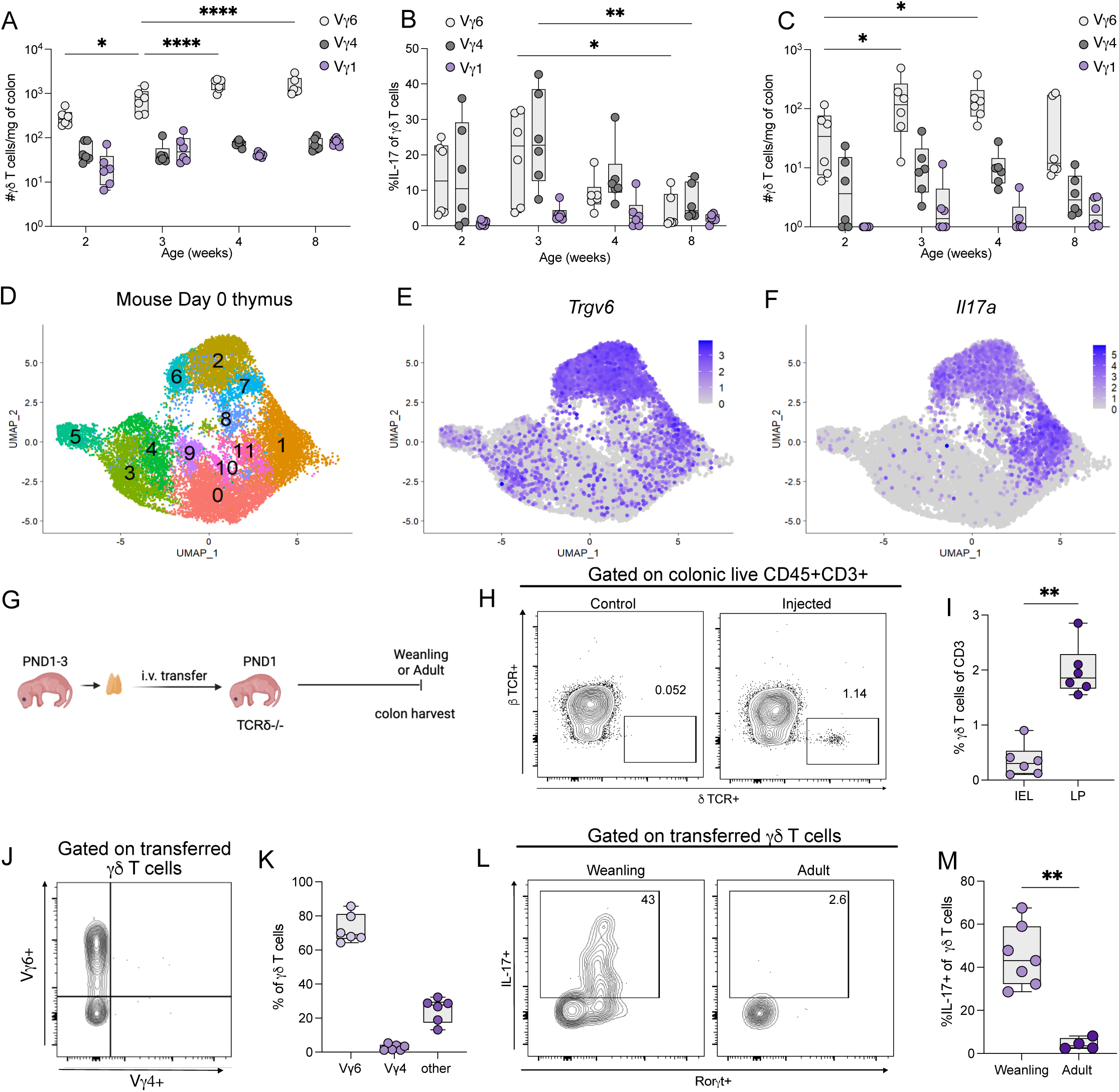
Fetal-derived. **V**γ**6+** γδ **T cells are activated in the early life colon. (A)** Total number of γδ T cells expressing Vγ6, Vγ4 or Vγ1 in the colon LP at indicated ages. **(B-C)** Proportion (B) and number (C) of IL-17+ γδ T cells expressing Vγ6, Vγ4 or Vγ1 in the colon LP of 2-, 3-, 4-, or 8-week-old mice. **(D-F)** UMAP of all cells (D), *Trgv6* (E) and *Il17a* (F) expression on total thymocytes from day 0 mouse **(G-I)** Mouse model (G), representative flow plot (H) and percent of γδ T cells (I) in the colon LP vs IEL of 3 week old mice following adoptive transfer of WT γδ T cells in 1-day old TCRδ-/-mice. **(J, K)** Representative flow plot (H) and percent of Vγ6+ and Vγ4+ γδ T cells in colon of mice that received γδ T cells. **(L, M)** Representative flow plot (L) and percent (M) of IL-17+ γδ T cells in colon LP of weanling and adult mice following adoptive transfer of γδ T cells. (A, B, C, I, K and M) Each dot represents cells from an individual mouse. Data are represented as mean with SD. Graphs are pooled data from at least two independent experiments with atleast n = 3 in each experiment. (H, J and L) Contour plots are representative at least two independent experiments. Numbers indicate frequency of cells within each quadrant or box. (A-C) Data were analyzed using 2-way ANOVA test or (I, M) Mann-Whitney non-parametric unpaired T test (∗p < 0.05, ∗∗p < 0.01, ∗∗∗p < 0.001, ∗∗∗∗p <0.0001).

### Early life γδ17 T cell activation is IL-1R-dependent

To examine the multicellular immune network that may shape colonic γδ T cell activation in weanlings, we returned to our single-cell sequencing dataset. In addition to γδ T cells, we found several age-dependent changes, including an enrichment of innate immune cells such as monocytes and innate lymphoid cells (ILCs) in weanling mice, compared to a greater proportion of adaptive immune cells (CD4+ and CD8+ αβ T cells, B cells and plasma cells) in the adult colon (Figure 3A). Unexpectedly, we observed a significant accumulation of neutrophils and monocytes in the weanling colon, a finding we validated by flow cytometry (Figure 3B, C). Since neutrophil recruitment to the gut has been shown to be dependent on IL-17 signaling in various contexts^20,37^ and colonic γδ T cells from the weanling colon highly expressed *Ccl5*, a monocyte chemokine, we hypothesized that IL-17, or γδ T cells more generally, would be similarly required in early life. However, colons from weanling TCRδ-/-mice and IL-17A-/-mice had similar numbers of neutrophils as littermate controls (Figure 3D, E; Figure S2A, B). Similarly, administration of anti-IL-17A antibody blockade from 2 to 3 weeks did not significantly change neutrophil or monocyte recruitment into the colon (Figure S2C, D). We therefore hypothesized that myeloid-lineage cells may be providing signals necessary for γδ17 T cell activation. To this end, we analyzed the top expressed genes in these populations in the weanling colon after removal of mitochondrial and ribosomal genes (Figure S2E, F). We found several genes such as *S100a6, S100a8, S100a9* highly expressed by both myeloid subsets, suggesting involvement in an early life antimicrobial response. We also found elevated *Il1b* expression elevated in both monocytes and neutrophils at 3 weeks of age (Figure 3F; Figure S2E, F), which has been previously shown to promote IL-17 production by γδ T cells^20^. Although neutrophils and monocytes expressed the greatest amount of *Il1b* mRNA, other myeloid cells such as dendritic cells and macrophages also expressed *Il1b* in the weanling colon (Figure S2G). Accordingly, we detected elevated levels of *Il1b* transcripts in colon tissue from 2-and 3-week-old mice that decreased with age (Figure 3G). Using flow cytometry, we also detected an increase in pro-IL-1β-producing CD11b+ myeloid cells at 3 weeks, compared to adult mice (Figure 3H, I). Similar results for mature IL-1β was confirmed from colon lysates (Figure 3J).

**Figure 3.**
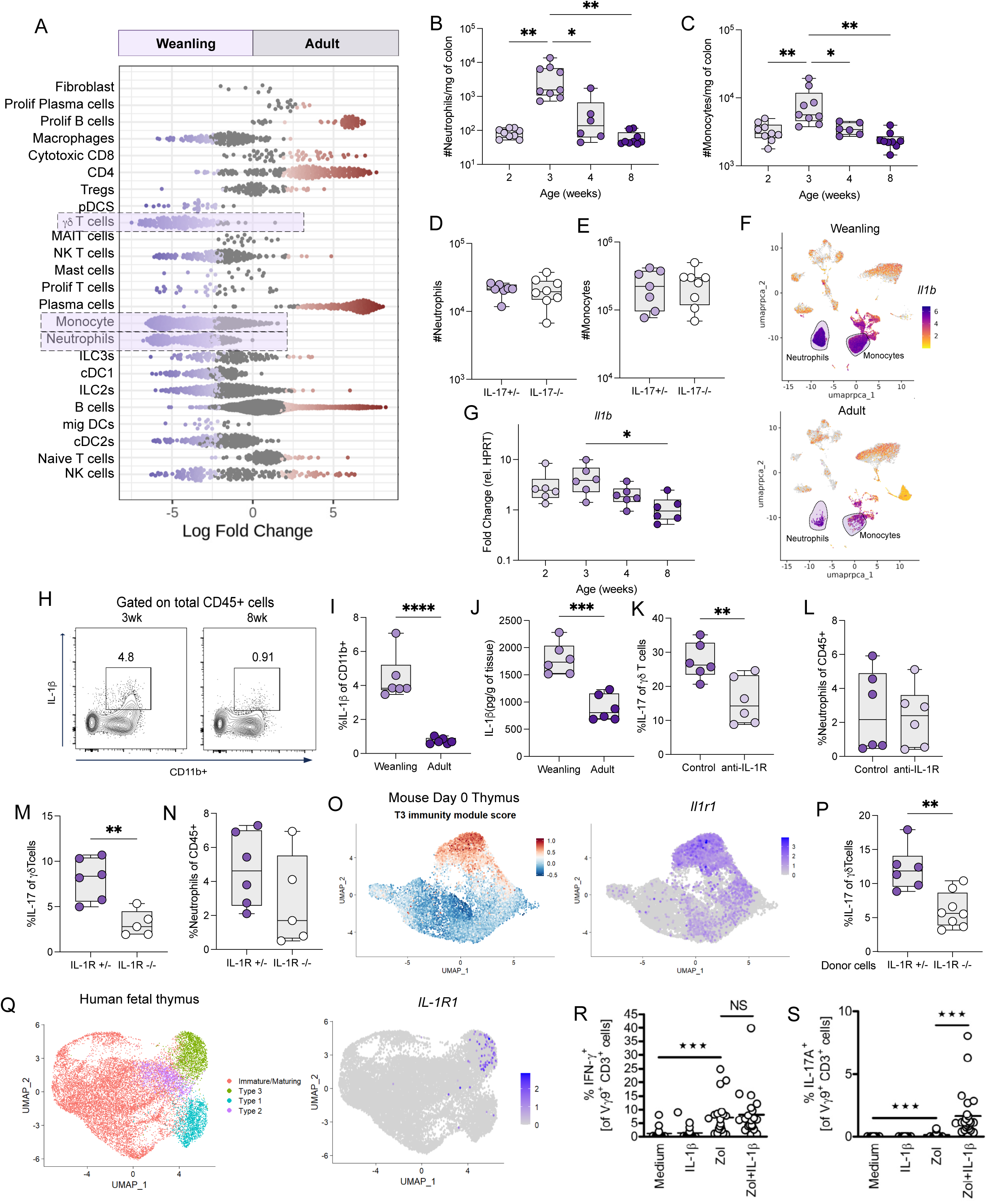
**The early life IL-17 response is dependent on IL-1**β **produced by infiltrating myeloid cells. (A)** Differential abundance using MiloR analysis of single cell sequencing of total CD45+ cells of weanlings and adults. **(B, C)** Total number of neutrophils(B) and monocytes (C) in the colon normalized to weight of the tissue of mice at indicated ages. **(D, E)** Total number of neutrophils (D) and monocytes (E) in the colon lamina propria of 3-week old littermate IL-17+/-and IL17-/-mice. **(F)** UMAP of *Il1b* expression in weanling and adult colon CD45+ cells. **(G)** Expression of *Il1b* in total colon of mice at indicated ages. **(H, I)** Representative flow plot (H) and percent (I) of pro-IL-1β+ myeloid cells in the LP of weanling and adult colons. **(J)** ELISA quantification of IL-1β in the supernatant of colon explants of weanling and adult mice, normalized to weight of tissue. **(K, L)** Percent of IL-17+ γδ T cells (K) and percent of neutrophils (L) in colonic LP of control mice and mice treated with anti-IL-1R in vivo antibody from 2-3 weeks. **(M, N)** Percent of IL-17+ γδ T cells (M) and percent of neutrophils (N) in colonic LP of littermate IL-1R+/-and IL-1R-/-at 3 weeks of age. **(O)** UMAP of type 3 immunity score and *Il1r1* expression on total thymocytes from day 0 mouse **(P)** Percent of IL-17+ γδ T cells in 3-week-old TCRδ-/-adoptively transferred with IL-1R+/-or IL-1R-/-thymus cells. **(Q)** UMAP of IL-1R1 expression in total CD45+ cells in human fetal thymus γδ T cells. **(R, S**) Percent of IFNγ (R) and IL-17 (S) of Vγ9+ γδ T cells in human cord blood following stimulation with IL-1β and zoledronate. (B, C, D, E, G, I-N, P, R, and S) Each dot represents cells from an individual mouse. Data are represented as mean with SD. Graphs are pooled data from at least two independent experiments with atleast n = 3 in each experiment. (H) Contour plots are representative at least two independent experiments. Numbers indicate frequency of cells within each quadrant or box. (B, C, G) Data were analyzed using Kruskal-Wallis ANOVA test or (D, E, I-N and P) Mann-Whitney non-parametric unpaired T test (∗p < 0.05, ∗∗p < 0.01, ∗∗∗p < 0.001, ∗∗∗∗p <0.0001).

To determine whether endogenous IL-1β plays a role in activating weanling γδ17 T cells, mice were treated with a neutralizing anti-IL-1R antibody from 2 to 3 weeks of age. IL-1R blockade reduced IL-17 production by γδ T cells, but no change was detectable in the number of neutrophils (Figure 3K, L), indicating again that the neutrophil recruitment is independent of IL-17 production. Similarly, IL-1R-/-weanlings had decreased IL-17 production but no change in neutrophil recruitment at 3 weeks of age compared to littermate controls (Figure 3M, N). To test whether IL-1-dependent IL-17 production by γδ T cells is cell-intrinsic, we returned to our neonatal cell transfer approach. Notably, scRNA sequencing of neonatal thymocytes indicated that Vγ6+ γδ T cells (clustered as type 3 gd T cells based on *Il17a* and *Rorc* expression also highly expressed *il-1r1* mRNA (Figure 3O). Therefore, we transferred 1 day-old thymocytes from IL-1R+/-or IL-1R-/-mice into neonatal γδTCR-/-mice, effectively repopulating the colonic γδ T cell compartment with or without IL-1R expression. We found that IL-1R-/-γδ T cells produced significantly less IL-17 in the weanling colon than their heterozygote counterparts, indicating that IL-1R directly promotes IL-17 production by γδ T cells (Figure 3P). Consistent with our mouse studies, type 3 γδ T cells in the human fetal thymus, as identified by expression of *RORC, CCR6* and *IL-17* from a previously published single cell dataset^38^ also expressed high amounts of *IL1R1* (Figure 3Q). To determine whether IL-1R signals promote human γδ T cells to produce IL-17, γδ T cells isolated from cord blood were stimulated with IL-1β and zoledronate, the latter being a bisphosphonate that stimulates the Vγ9Vδ2 T cell receptor^39^. Unlike IFNγ production which was independent of IL-1β stimulation, IL-1β significantly increased the production of IL-17 in zoledronate-treated γδ T cells (Figure 3R, S). Collectively, these data demonstrate that IL-1β promotes IL-17 production by colonic γδ T cells in early life mouse and human γδ T cells.

### IL-10 extinguishes γδ T cell activation in a cell-extrinsic manner

As shown above, γδ17 T cell activation was transient, occurring only at 3 weeks of age. These results led us to investigate the mechanisms responsible for terminating IL-17 production in the adult colon. As previously mentioned, the T cell population in the weanling colon is drastically different to that of the adult colon (Figure 4A), where the latter shifts to have a greater proportion of conventional CD8+ and CD4+ αβ T cell populations such as Th17 cells and regulatory T cells (Treg). Treg clusters were compared to a previously published single-cell sequencing dataset of Tregs the colon defining a suppressive subset and a non-lymphoid tissue subset^40^. Interestingly, the adult colon had increased suppressive Tregs defined by elevated levels of *Il10* and *Lag3* mRNA (Figure 4B; Figure S2H, I). Consistently, IL-10 production has been shown to limit type 3 immune responses^41^. Using flow cytometry, we found that Rorγt+ Treg cells began to accumulate in the colon at 3 weeks of age, whereas there was no detectable change in the percent of Rorγt-Treg over time (Figure 4C-E). Indeed, the Il10 expressing Treg cluster was also enriched for *Rorc* mRNA (Figure S2J). In the adult colon, Rorγt+ Tregs were the main producers of IL-10 following *ex vivo* PMA and ionomycin stimulation (Figure 4F). Indeed, the total number of IL-10+ Rorγt+ Tregs significantly increased with age (Figure 4G). To test whether IL-10 plays a role in extinguishing γδ17 T cell activation, we administered a neutralizing anti-IL-10R antibody to mice from 3 weeks to 5 weeks of age. We found that blockade of IL-10 signaling sustained IL-17 production by γδ T cells compared to isotype control treated mice (Figure 4H). We also found increased neutrophil recruitment and IL-1β production in IL-10R neutralized mice (Figure 4I, J) suggesting regulation of γδ17 activation by IL-10 may occur at the level of recruited myeloid cells. Indeed, IL-10 stimulation *in vitro* did not directly limit IL-17 production by γδ T cells following IL-1β and IL-23 stimulation (Figure 4K, L). Instead, we found that IL-10 stimulation limited ATP + LPS-induced IL-1β production by colonic myeloid cells (Figure 4M, N). These data indicate that IL-10, likely produced by Rorγt+ Tregs limits IL-1β production which, in turn, downregulates γδ17 T cell activation. Notably, IL-10R blockade also significantly increased the number of IFNγ and IL-17 producing CD4+ T cells (Figure 4O, P) as well as colon thickness and tissue pathology (Figure 4Q, R). Together, these data suggest that accumulation of IL-10 producing Rorγt+ Tregs subsequent to γδ17 T cell activation terminates the type 3 immune response to prevent tissue pathology.

**Figure 4.**
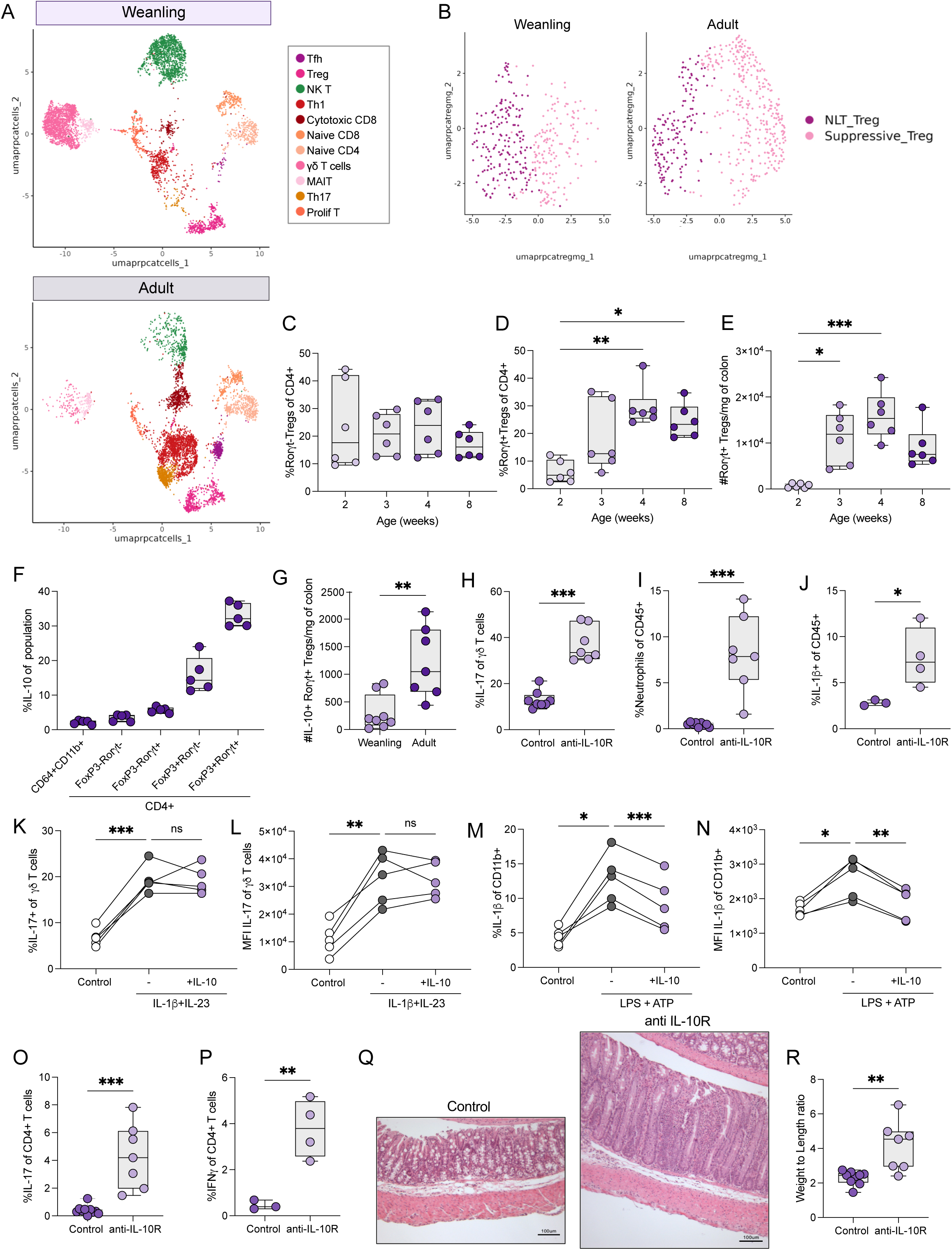
γδ**17 T cell activation is limited in adulthood by IL-10 producing Ror**γ**t+ regulatory T cells. (A)** UMAP of total CD3+ in the colon LP of weanling and adult mice. **(B)** UMAP of regulatory cells expressing non-lymphoid tissue regulatory T cell phenotype of suppressive phenotype in total Tregs. **(C, D)** Percent of Rorγt-(C) and Rorγt+ (D) Tregs of total CD4+ cells of the colons of mice at indicated ages. **(E)** Total number of Rorγt+ Tregs in the colons of 2-, 3-, 4-, and 8-week-old mice, normalized to weight of the tissue. **(F)** Percent IL-10+ cells of different immune cell subsets in the colon LP of adult mice following 4 hours of PMA and ionomycin stimulation. **(G)** Total number of IL-10+ Rorγt+ regulatory Tregs in the colon of weanling and adult mice, normalized to weight of tissue. **(H-J)** Percent of IL-17 by LP γδ T cells (H), percent of neutrophils (I) and IL-1β+ myeloid cells (J) in the colon of control or IL-10R in vivo antibody treated mice. **(K, L)** Percent (K) and MFI (L) of IL-17+ γδ T cells in ex vivo stimulation of total colon LP cells with IL-1β (10ng/ml) and IL-23 (10ng/ml), with or without IL-10 (10ng/ml). **(M, N)** Percent (M) and MFI (N) of IL-1β+ CD11b+ cells in ex vivo stimulation of total colon LP cells with ATP (3mM) and LPS (100ng/ml), with or without IL-10 (10ng/ml). **(O, P)** Percent of IL-17+ (O) and IFNγ+ (P) CD4+ T cells in the colon lamina propria of control mice and IL-10R in vivo antibody treated mice. **(Q, R)** H and E images (Q) and weight to length ratio (R) of colons of control mice and IL-10R in vivo antibody treated mice. (C-J, O, P, R) Each dot represents cells from an individual mouse or (K-N) each line represents a biological replicate of 3 mice combined. Data are represented as mean with SD. Graphs are pooled data from at least two independent experiments with atleast n = 3 in each experiment. (C-E) Data were analyzed using Kruskal-Wallis ANOVA test or (G-J, O, P, and R) Mann-Whitney non-parametric unpaired T test or (K-N) using the Wilcoxon nonparametric paired t test (∗p < 0.05, ∗∗p < 0.01, ∗∗∗p < 0.001, ∗∗∗∗p <0.0001).

### γδ17 T cell activation is microbiota-dependent and-specific

Previous studies have indicated that intestinal γδ17 T cell function is shaped by the gut microbiota^27,28^. However, its impact on γδ17 T cell activation during early life has not been investigated. To examine this possibility, we measured γδ17 T cell responses in 3 week-old specific pathogen-free (SPF) mice and germ-free mice (GF). GF mice had a small, but significant decrease in the total number of γδ T cells in the weanling colon compared to SPF controls, possibly due to decreased proliferation (Figure S3A, B). Nevertheless, the majority of colonic γδ T cells retained expression of Rorγt in the absence of a microbiota (Figure 5A) and were able to produce IL-17 upon in vitro stimulation (Figure 5B). Actually, γδ T cells from GF mice were able to produce slightly but significantly more IL-17 following in vitro IL-1β and IL-23 stimulation, corroborating data from a previous study^28^. In contrast, ex vivo IL-17 and IL-22 production by colonic γδ T cells was abrogated in GF weanlings (Figure 5C-E), suggesting that in vivo activation of γδ T cells during early life requires microbial-derived factors. To determine whether microbiota-mediated IL-17 production by γδ T cells occurred in an age-dependent manner, we reconstituted adult GF mice with SPF feces and measured γδ17 T cell activation over time (Figure 5F). We found that, by 7 days post-colonization of adult mice, there was a significant increase in proliferation, IL-17 and IL-22 production by γδ T cells (Figure 5G, H; Figure S3C), indicating γδ17 T cell activation by the gut microbiota is not strictly age-dependent.

**Figure 5.**
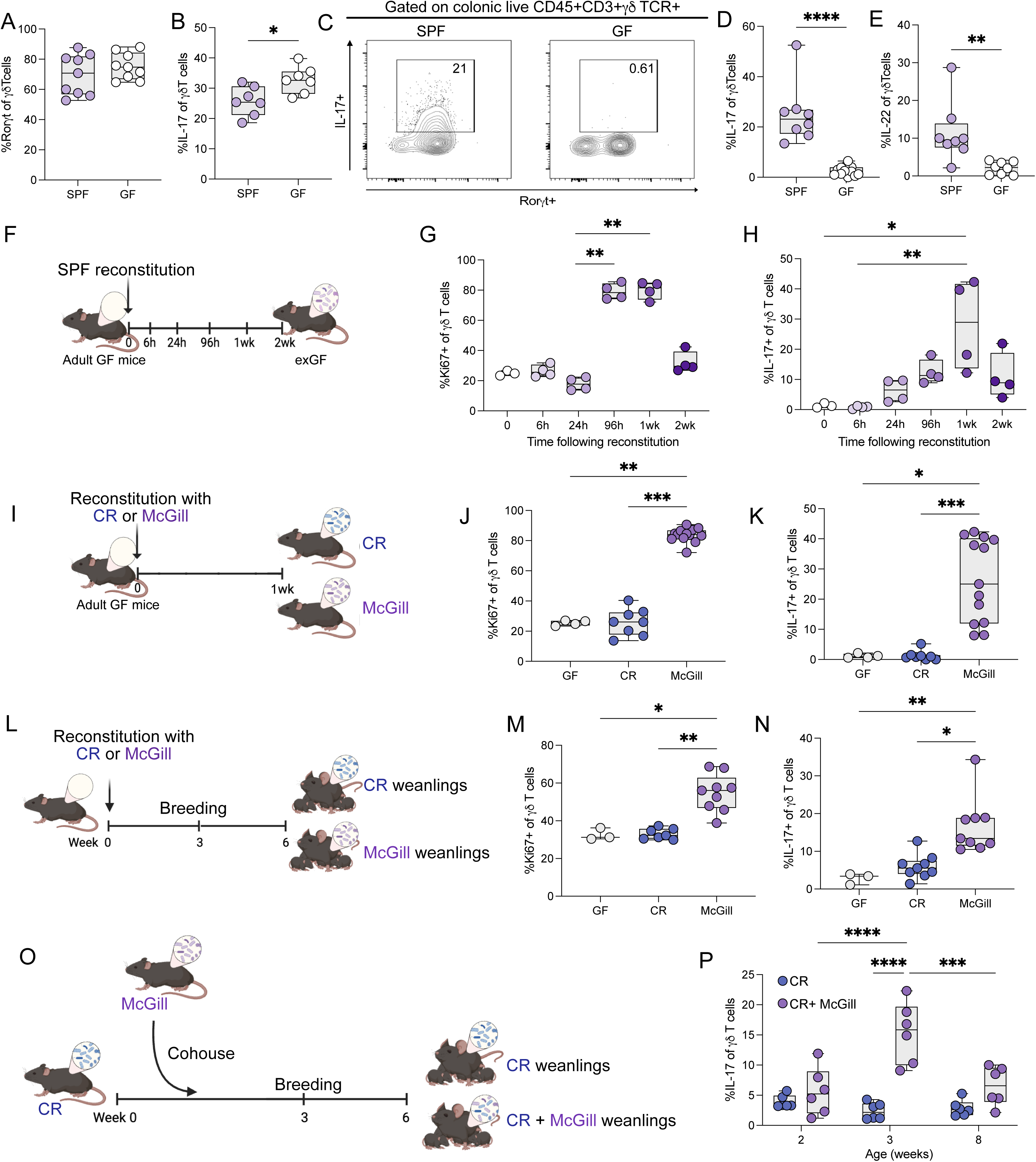
Early life IL-17 responses are microbiota-dependent and-specific. **(A)** Percent of Rorγt+ γδ T cells in weanling SPF and GF mice. **(B)** Percent of IL-17+ colonic LP γδ T cells following 4-hour stimulation with 10ng/ml IL-1β and 10ng/ml IL-23. **(C-E)** Representative flow plot (C) and percent (D) of IL-17+ γδ T cells and percent of IL-22+ γδ T cells (E) in the colon of SPF and GF mice. **(F-H)** Mouse model (F), percent of Ki67+ (G) and IL-17+ (H) γδ T cells in the colon following GF reconstitution at indicated times n = 4. **(I-K)** Mouse model (I), percent of Ki67+ (J) and IL-17+ (K) γδ T cells in the colon of adult GF mice reconstituted with CR or McGill SPF microbiotas. **(L-N)** Mouse model (L), percent of Ki67+ (M) and IL-17+ (N) γδ T cells in the colon of 3-week old pups with CR or McGill SPF microbiotas. **(O, P)** Mouse model (O) and percent IL-17+ γδ T cells (P) in the colon of mice with CR microbiota or CR+ McGill microbiota at indicated ages. (A,B, D, E, G, H, J, K, M, N, P) Each dot represents cells from an individual mouse or (K-N) each line represents a biological replicate of 3 mice combined. Data are represented as mean with SD. Graphs are pooled data from at least two independent experiments with atleast n = 3 in each experiment, unless otherwise indicated. (C) Contour plots are representative at least two independent experiments. Numbers indicate frequency of cells within each quadrant or box. (G, H, J, J, M, N) Data were analyzed using Kruskal-Wallis ANOVA test or (A, B, D, E) Mann-Whitney non-parametric unpaired T test or (P) using 2-way ANOVA test (∗p < 0.05, ∗∗p < 0.01, ∗∗∗p < 0.001, ∗∗∗∗p <0.0001).

To determine whether the specific composition of the microbiota, not just the microbiota per se, drives the colonic γδ17 T cell response, we compared adult mice colonized with SPF microbiota from our facility (denoted as McGill) to adult mice colonized with SPF microbiota obtained from mice bred at Charles River laboratories (denoted as CR) (Figure 5I). Remarkably, we found that only γδ17 T cells from McGill microbiota-exposed, but not CR microbiota-exposed mice, produced IL-17, IL-22 and increased proliferation (Figure 5J, K, Figure S3D). To test whether this also happens in early life, we bred adult GF mice reconstituted with either McGill or CR feces and performed flow cytometry on the colons of their 3-week-old pups (Figure 5L). Similar to the microbiota transfers in adult mice, IL-17, IL-22 and proliferation was induced in γδ17 T cells from 3-week-old colonized with McGill, but not CR microbiota (Figure 5M, N, Figure S3E). To determine whether the McGill microbiota exerted dominant effects over the CR microbiota in terms of the γδ17 T cell response, we cohoused female mice harboring McGill and CR microbiota for 3 weeks prior to breeding with CR males (Figure 5O). The progeny of cohoused dams had increased γδ17 T cell activation at 3 weeks compared to progeny of the original CR dams (Figure 5P).

To examine global differences in the colonic immune compartments between McGill-and CR-colonized 3-week-old mice, we performed a new scRNAseq experiment (Figure S3F). As expected, these results confirmed the greater proportion of γδ T cells in the colon of McGill vs CR microbiota mice as well as the increased presence of γδ T cells, neutrophils and monocytes in the former (Figure S3G). We also found increased IL-1β producing myeloid cells and increased neutrophil recruitment in McGill microbiota-colonized mice compared to their CR-colonized counterparts (Figure S3H, I). These data indicate that activation of weanling γδ17 T cells in the colon is specific to the composition of the gut microbiota and that this microbiota can be transferred vertically and horizontally.

### Intergenerational transfer of *C. difficile* drives γδ17 T cell activation during early life

To determine which microbe(s) is responsible for the activation of γδ17 T cells in early life, we performed 16S rDNA sequencing on fecal samples of McGill and CR mice. The McGill microbiota had a lower overall alpha diversity indicated by Shannon index values (Figure 6A). Using Bray-Curtis distance to assess β-diversity, we found significant differences in community composition (Figure 6B). To identify specific bacteria that differ between groups, we performed differential abundance analysis using LefSE^42^. We found changes in the abundance of several bacterial species between McGill and CR microbiota including a significant enrichment of probiotic species (e.g. *Bifidobacteria* and *Lactobacillus* spp.) as well as known pathobionts such as *Helicobacter typhlonius*, *Escherichia coli*, and *Clostridioles difficile* in the McGill microbiome compared to the CR microbiome (Figure 6C). Since intentional neonatal infection with *C.difficile* has been previously demonstrated to drive IL-17 production by γδ T cells^43^, we focused our attention on this pathobiont (Figure 6D). We confirmed that *C.difficile* 16S rRNA was present in the McGill microbiota, yet undetectable in CR microbiota by species-specific qPCR (Figure 6E). We next confirmed the presence of live *C.difficile* by plating McGill and CR feces on *C.difficile*-specific CDBA plates; an average of 1 x 10^6^ CFU of *C.difficile* per gram of feces in McGill mice was detected, but no growth in CR fecal cultures (Figure 6F, G). Furthermore, *C.difficile* in McGill mice actively expressed toxins *TcdA* and *TcdB in vivo* (Figure 6H, I). While *C.difficile* is an important cause of adult diarrheal disease in hospital-acquired infections^44^, our weanling mice did not exhibit any sickness behavior. This presentation mimics the clinical observation that up to a third of babies under six months of age are asymptomatically colonized with *C. difficile*, but decreases over time^45^.

**Figure 6:**
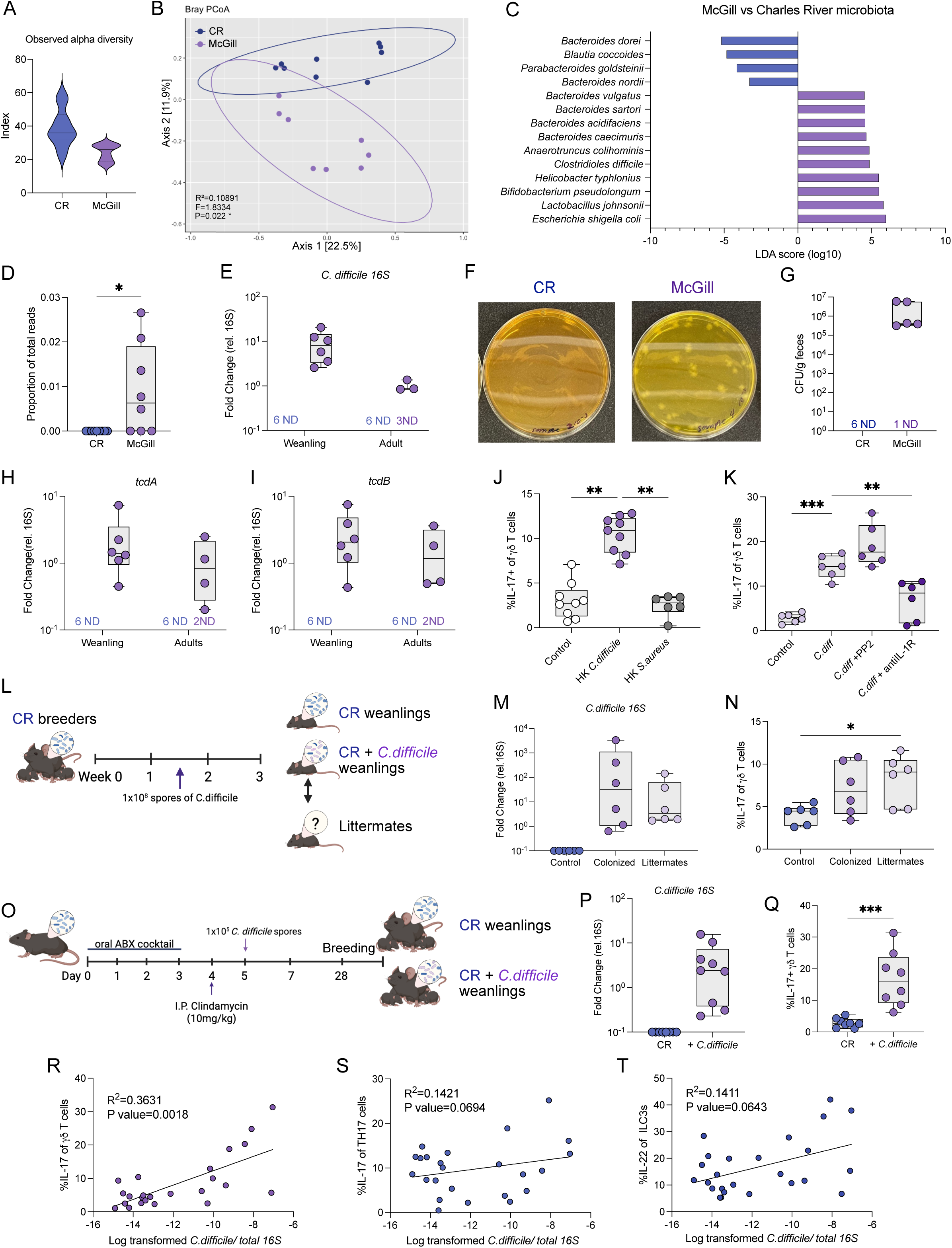
E**a**rly **life** γδ**17 T cell activation occurs following passive acquisition of *C. difficile*. (A-D)** Shannon Index (A), Bray-Curtis PCoA ordination plot (B), Differential abundance of ASVs associated to known bacteria using LefSE with threshold set at p>0.05 (C) and proportion of *C. difficile* 16S sequences (D) in 16S sequencing of CR and McGill microbiotas. **(E)** Expression of *C. difficile* 16S relative to total 16S in the feces of weanling and adult CR and McGill mice. **(F)** Representative images of CDBA plates with CR or McGill feces. **(G)** CFU of *C. difficile* found in CR and McGill feces. **(H-I)** Expression of *Tcda* (H) and *Tcdb* (I) in feces of weanling and adult CR and McGill mice. **(J)** Percent of IL-17+ γδ T cells following 24-hour stimulation of adult colon cells with heat-killed(HK) *C. difficile* and HK *S. aureus.* **(K)** Percent of IL-17+ γδ T cells following 24-hour stimulation of adult colon cells with HK *C. difficile* with PP2 (5μm) or anti-IL-1R (20ng/ml). **(L-N)** Mouse model (L), *C. difficile* 16S expression (M), and percent of IL-17+ γδ T cells (N) in the colon of CR pups colonized with *C. difficile* spores and their littermates. **(O-Q)** Mouse model (O), *C. difficile* 16S expression (P), and percent of IL-17+ γδ T cells (Q) in the colon of CR control pups and pups of *C. difficile* colonized breeders. **(R-T)** Correlation of *C. difficile* 16S rDNA and percent of IL-17+ γδ T cells (R), IL-17+ Th17 cells (S) and IL-22+ ILC3s (T) in weanlings. (D, E, G, H-K, M, N, P-T) Each dot represents cells from an individual mouse or (K-N) each line represents a biological replicate of 3 mice combined. Data are represented as mean with SD. Graphs are pooled data from at least two independent experiments with at least n = 3 in each experiment. (J, K, M, N) Data were analyzed using Kruskal-Wallis ANOVA test or (P, Q) Mann-Whitney non-parametric unpaired T test or (R-T) using Pearson correlation test (∗p < 0.05, ∗∗p < 0.01, ∗∗∗p < 0.001, ∗∗∗∗p <0.0001).

We next sequenced the *C.difficile* strain found in our McGill microbiota mice to determine the strain and potential origin of aquisition. *C. difficile* is classically categorized by clades based on multilocus sequence typing^46^. Hospital-acquired strains typically belong to clade 2 whereas lab strains are typically clade 1 due to their less virulent nature. We performed full-length 16S rRNA nanopore sequencing of our unknown strain and commonly used lab strain VPI1063 as a control. Our unknown strain was closely related to clade 2 strains of *C.difficile*, including the common Quebec outbreak strain RB027, demonstrated by very high average nucleotide identity scores (Figure S4A, B). This suggests that colonization of our mice with *C.difficile* most likely occurred by an external source, and not through experimental contamination.

We next tested if the presence of *C.difficile* in the McGill microbiota can induce γδ17 activation in early life. *In vitro* stimulation of total colonic LP cells with heat-killed *C.difficile*, isolated from McGill microbiota, but not another gram-positive bacteria *S. aureus*, induced IL-17 production by colonic γδ T cells (Figure 6J). We also found elevated IL-1β in the supernatant of *C.difficile* stimulated LP cells (Figure S4C). Indeed, blocking IL-1R signalling during *in vitro* stimulation abrogated IL-17 production by colonic LP γδ T cells (Figure 6K). Notably, blocking TCR signalling using LCK-inhibitor PP2 did not affect IL-17 production by LP γδ T cells, suggesting that γδ17 T cell activation in this setting is independent of TCR engagement. To test whether *C. difficile* drives γδ17 T cell activation *in vivo*, we gavaged half a litter of pups at 10 days post-birth with 1×10^8^ CFU of *C.difficile* spores, leaving the remaining pups untouched (Figure 6L). At weaning, we detected *C. difficile* in pups that were directly gavaged with *C.difficile* as well as littermate siblings of these pups that were not gavaged, indicating that early life horizontal transfer of *C. difficile* occurred (Figure 6M). Importantly, we found elevated number of γδ T cells and increased proportion of IL-17 producing γδ T cells in mice reconstituted with *C.difficile* (Figure 6N; Figure S4D). Interestingly, IL-22 production by γδ T cells was unchanged following *C.difficile* colonization, indicating that the requirements for microbiota-mediated IL-17 and IL-22 production by γδ T cells may be distinct (Figure S4E). Next, we tested whether colonization of the breeders would lead to a similar activation of γδ17 T cells in the progeny. As *C.difficile* does not efficiently colonize adult mice harboring a mature microbiota, we treated adult CR microbiota-colonized mice with a short antibiotic regimen prior to gavaging them with 1×10^5^ *C.difficile* spores (Figure 6O). After confirming colonization, mice were bred to examine the γδ17 T cell immune response in the progeny. We confirmed that pups from *C.difficile*-colonized breeders also had *C.difficile* in their microbiota indicating that vertical transfer occurred (Figure 6P). Accordingly, we found that *C.difficile* induced both proliferation and IL-17 production by colonic γδ T cells from weanling mice (Figure 6Q, Figure S4F). Again, IL-22 production by γδ T cells was unchanged (Figure S4G). Neutrophil recruitment and IL-1β production by CD11b+ myeloid cells was also significantly increased in the colonic LP of pups colonized with *C.difficile* colonized to non-colonized controls (Figure S4H, I). Interestingly, we found that the amount of *C. difficile* 16S rRNA detected in the feces positively correlated with IL-17 production by γδ T cells (Figure 6R), but did not significantly correlate with either IL-17 production by Th17 cells or IL-22 production by ILC3s (Figure 6S, T). Together, these data indicate that *C.difficile* can be transferred between mice, both horizontally and vertically, and passive acquisition of *C.difficile* from dams to pups induces IL-17 by γδ T cells.

### Early life IL-17 production promotes intestinal barrier integrity and controls *C. difficile* persistence

During the course of our studies, we also observed a microbiota-dependent accumulation of Vγ6+ Rorγt+ γδ T cells in the epithelial fraction of the colon at 3 weeks of age (S4J, K), an area normally populated exclusively by Vγ7+ γδ T cells that do not express Rorγt or produce IL-17^47–49^. Furthermore, we detected elevated intercrypt bacterial DNA in the colon of weanlings harbouring a McGill microbiota, potentially due to epithelial damage caused by *C.difficile* toxins (Figure 7A, B). Furthermore, McGill weanling mice had increased levels of TLR4 ligands in the serum, measured using a reporter cell line, which indicates further bacterial dislocation into the systemic circulation (Figure 7C). Combining these data with previous results showing that *C. difficile* can compromise epithelial barrier integrity and increase dissemination of commensal bacterial species to the systemic circulation^50^, we hypothesized that LP γδ17 T cells accumulate in the intraepithelial to directly influence epithelial function. In support of our hypothesis, there was a significant increase in expression of the tight junction protein, ZO-1, in the epithelium of McGill weanlings compared to CR microbiota-colonized mice (Figure 7D, E). Since we did not observe any difference in overt sickness behavior between groups despite changes at the tissue and serum levels, we hypothesize that the host adapts to tolerate a *C. difficile*-containing microbiota during early life by modulating immune-epithelial interactions.

**Figure 7.**
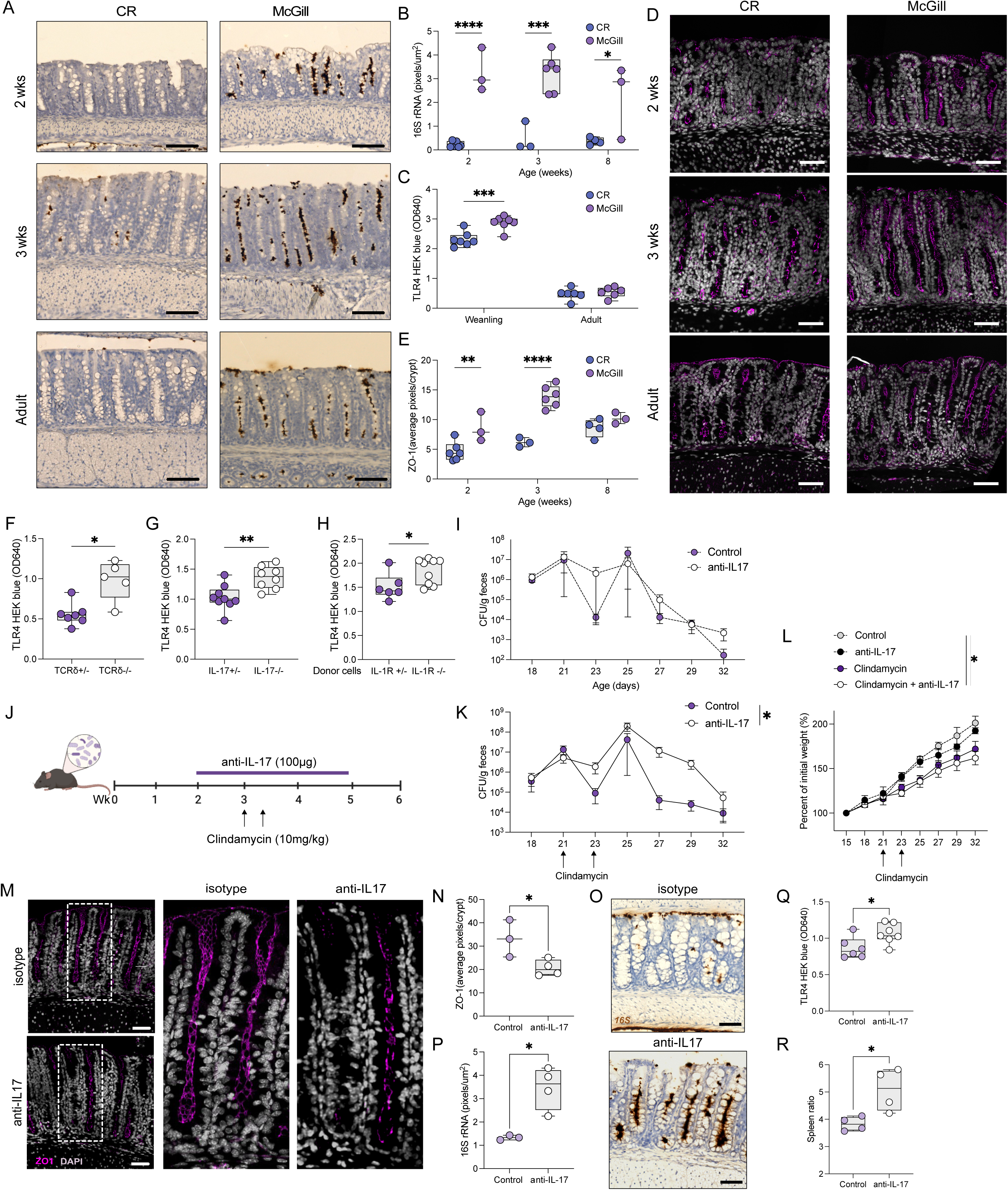
Early life IL-17 limits bacterial translocation by strengthening the epithelial barrier. (A, **B)** RNAScope images (A) and quantification (B) of 16S transcripts in CR and McGill microbiota mice at indicated ages. **(C)** OD640 of TLR4 HEK cell line following 24-hour incubation with serum from weanling and adult CR and McGill mice. **(D, E)** Confocal image of ZO-1 staining (D) and quantification (E) in CR and McGill mice at indicated ages. **(F-H)** OD640 of TLR4 HEK cell line following 24-hour incubation with serum from littermate δTCR+/-and δTCR-/-mice (F), littermate IL-17+/-and IL-17-/-mice (G)and δTCR-/-that received IL-1R+/-or IL-1R-/-γδ T cells adoptive transfer (H). **(I)** CFU of *C. difficile* in control and anti-IL-17 (6 doses of 100mg/ml every 3 days) starting at 2 weeks. **(J-L)** Mouse model (J), *C. difficile* CFU in feces (K) and weight (L) of McGill mice treated with anti-IL-17 and two doses of clindamycin (10mg/kg) at weaning. **(M, N)** Confocal image of ZO-1 staining (M) and quantification (N) in control and anti-IL-17 treated mice. **(O, P)** RNAScope images (O) and quantification (P) of 16S transcripts in control and anti-IL-17 treated mice. **(Q)** OD640 of TLR4 HEK cell line following 24-hour incubation with serum from control and anti-IL-17 treated mice (**R**) Spleen size, normalized to weight of mouse in control and anti-IL-17 treated mice. (B, C, E, F-H, N, P, Q, and R) Each dot represents cells from an individual mouse. Data are represented as mean with SD. Graphs are pooled data from at least two independent experiments with atleast n = 3 in each experiment. (B, C, E, K, L) Data were analyzed using two-way ANOVA test or (F-H, N, P, Q and R) Mann-Whitney non-parametric unpaired T test (∗p < 0.05, ∗∗p < 0.01, ∗∗∗p < 0.001, ∗∗∗∗p <0.0001). Scale bar, 50μm.

To directly test whether this adaptation was influenced by the presence of γδ T cells and/or IL-17A, we tested serum samples in our TLR4 assay isolated from weanling *Tcrd-/-* and *Il17a*-/-mice with a McGill microbiota. Indeed, both mouse lines had increased amounts of TLR4 ligands in the serum compared to littermate controls at weaning age (Figure 7F, G). Moreover, *Tcrd-/-* mice reconstituted with IL-1R-sufficient neonatal thymocytes had a significantly lower amount of TLR4 ligands in their serum compared to mice that received IL-1R-deficient neonatal thymocytes (Figure 7H).

Since *C. difficile* CFUs positively correlated with IL-17 production by γδ T cells (Figure 6Q) and that bacterial burden was greater in weanling mice compared to the same mice in adulthood (Figure 6E), we hypothesized that IL-17A production in early life not only contributes to epithelial barrier integrity, but also promotes clearance of *C.difficile* to prevent persistence of this pathobiont. To test this hypothesis, we treated weanlings from McGill microbiota-colonized mice with anti-IL-17A to directly determine the role of IL-17 on pathobiont growth and maintenance. Under these conditions, IL-17 blockade alone did not impact the number of *C. difficile* fecal CFU (Figure 7I) indicating, as previously described^51^, that a replete gut microbiota efficiently controls growth of this pathobiont. We then performed a similar experiment except mice were given two doses of Clindamycin at weaning to simulate antibiotic-induced dysbiosis that can lead to disease in hospital-acquired *C. difficile* infection^52^ (Figure 7J). In this case, IL-17A blockade led to delayed clearance of *C.difficile* and failure to gain as much weight with age compared to isotype control-treated mice (Figure 7K, L). Consistently, anti-IL-17A treatment also led to a decrease in epithelial ZO-1 expression and increased detection of intercrypt bacterial DNA (Figure 7M-P). Moreover, anti-IL-17 treatment in the context of clindamycin led to an increase in serum TLR4 ligand detection and larger spleens compared to isotype-treated mice (Figure 7Q, R).

## DISCUSSION

By investigating the dynamics of the immune response within the colonic microenvironment at different ages of mice, we describe a multicellular network that acts in a rapid yet restrained manner during early life to promote epithelial barrier integrity, limit pathobiont growth and prevent immunopathology. Specifically, we observed that γδ17 T cells selectively expanded and produced type 3 cytokines including IL-17 and IL-22 at the age of weaning. A previous study also reported a transient intestinal cytokine response at the same age, characterized by production of IFNγ and TNF in the ileum, a phenomenon they termed the weaning reaction^34^. Consistent with this study, we also observed transient production of IFNγ by ileal CD4+ T cells at weaning age. Both studies demonstrated a critical role for the microbiota in driving the immune response at this time as well as a role for Rorγt+ Tregs for prevent immunopathological responses with age. These data are consistent with a longitudinal study in humans where increased detection of plasma and fecal IL-17A, coincident with increased proportion of γδ T cells with a gut-homing phenotype^53^, prior to or at the time of weaning closely tracked with a specific gut microbiota composition at the same age^32^. Moreover, this study demonstrated an inverse relationship between serum IL-17 and IFNγ production. Interestingly, this study also showed an increase in the frequency of circulating memory Foxp3+ Tregs immediately following inflammatory cytokine detection^32^. Taken together, we propose that the weaning reaction and its associated mechanism of regulation is a conserved phenomenon across intestinal microenvironments, yet the quality of the reaction depends on the microbiota to which it is responding.

In our study, early life colonization with a microbiota harboring *C. difficile* induced IL-1β production by diverse myeloid cell populations necessary to activate γδ17 T cells. These data are consistent with a previous study demonstrating that the gut microbiota increases IL-1R expression on γδ17 T cells^27^. We build on these results by showing that IL-1R is highly expressed on mouse and human fetal thymic γδ17 T cells and, using an adoptive transfer approach, demonstrate that IL-1R signaling is important for microbiota-mediated activation of γδ17 T cells in vivo. These data are in agreement with a study showing that neonatal human Vγ9Vδ2+ γδ T cells are predisposed to producing IL-17 compared to their adult counterparts^54^. These data, combined with a known role for IL-1 in activating γδ17 T cells at diverse barrier sites^16,27,55^, suggest that IL-1R is likely part of a larger γδ17 T cell transcriptional program imprinted during thymic development and retained in early life^4^.

Even though *C. difficile* remains one of the most prevalent causes of antibiotic-induced infectious colitis in North American adults^56^, the detection of *C. difficile* in asymptomatic is very common^57^. In fact, *C.difficile* was first described in 1935 as part of the normal flora of newborn children^58^ and a more recent meta-analysis summarizing results from over a hundred clinical studies showed asymptomatic colonization of more than a third of children, with a peak at 2-4 months of age^45^. Similarly, the acquisition of *C. difficile* by pups in our mouse colony also occurred without overt symptoms, as observed in a previous study in mice^59^, mimicking the clinical scenario of asymptomatic carriage described above. Thus, we took advantage of this opportunity to study the early life immune response and its impact barrier integrity during *C. difficile* colonization. While we observed a robust γδ17 T cell response that limited pathobiont growth and enhanced barrier defense, we did not observe severe disease, even in the presence of antibiotics or when IL-17A was neutralized. This contrasts with a previous study showing that γδ17 T cells prevent against severe disease in neonates^60^. These differing results may be due to several reasons. First, we examined mice that were colonized with this pathobiont from birth. These conditions may enable the developing microbiota to adapt to the presence of this pathobiont and limit its virulence^61^. Second, even though the *C. difficile* strain that we investigated is genetically similar to hypervirulent strains and expresses tissue-damaging toxins, its passage in our animal colony over multiple generations may have altered its metabolic niche and dampened its virulence^62^. In addition, we treated mice already colonized with *C. difficile* with a limited regimen of clindamycin in an attempt to approximate the clinical setting where this single antibiotic is commonly associated with overt *C. difficile* infection^63^. However, this experimental design differs from most preclinical studies in which animals are treated with a cocktail of broad-spectrum antibiotics prior to a bolus bacterial challenge^50,60,64^. As such, microbiota-mediated control of *C. difficile* growth may not be as compromised in our experimental design as previous studies. Another possibility is that other host-derived pathways may partially compensate for the absence of IL-17. Indeed, IL-22 has been shown to contribute to protection against *C. difficile* in a complementary manner to IL-17 by limiting systemic dissemination of commensal bacteria^50^. Although a mechanism was not described, IFNγ has also been shown to control *C. difficile* growth and limit intestinal damage^64^. None of these potential explanations are mutually exclusive and deserve further investigation.

### Limitations of the study

While our study examined how γδ17 T cells contribute to colonic host defense during the early life period, the long-term implications of this response remain to be determined. While we described the microbiota-dependent accumulation of gd17 T cells in the intraepithelial layer during the weaning period, the signals that drive this change in localization and whether this migration is critical for their effects on barrier integrity are unclear. In addition, the microbiota-derived signals that activate myeloid cells to produce IL-1β in the presence of *C. difficile* have yet to be described. Finally, additional studies will need to be performed to determine whether *C. difficile* is unique in its ability to activate γδ17 T cells during early life or whether other pathobionts known to compromise the epithelial barrier can drive this type 3 immune response in a similar manner.

## EXPERIMENTAL MODEL AND STUDY PARTICIPANT DETAILS

### Mice

All experimental procedures were performed in accordance with the McGill University Health Centre Research Institute Animal Resource Division with approved use protocol #7977. Wildtype McGill and wildtype CR mice (C57BL/6, No. 027) were both obtained originally from Charles River. Il17a^cre^ knock-in mice (C57BL/6, No.035717) were obtained from Kathy McCoy (University of Calgary) and bred to generate Il17aCre/+ and Il17aCre/Cre pups used for experiments. TCRδ-/-mice (C57BL/6, No.002120) were obtained from Jackson laboratories. IL-1r1-/-mice (C57BL/6, No.003245) were obtained from Jackson laboratories. Germ free mice on a C57BL/6 background were purchased from the International Microbiome Centre at the University of Calgary and bred at the McGill Centre for Microbiome Research gnotobiotic animal research platform. All mice were backcrossed to the C57BL/6 background and transgenic lines were bred in house. Mice were housed in groups (4-5 mice per cage) with *ad libidum* access to standard chow and water. Animals were housed at 22C with a 12h cycle of lights on (7am) and lights off (7pm). For all mouse lines, littermates of the same sex were randomly assigned to experimental groups. Sex had no effect on the experimental outputs, so all combined data is expressed as a combination of the sexes. Health of the mice was monitored every 2-3 days or daily.

### Antibody treatment

For neutralization of IL-17a activity, mice were treated i.p. with 100 μg of anti-IL-17A antibody (clone 17F3, BioXcell) starting on day 14 post-birth and every second day thereafter until euthanized. For IL-10R blocking experiments, mice were treated i.p. with 200 μg of in-vivo anti-IL-10R antibody (clone 1B1.3A, BioXcell) starting at 3 weeks and every other day until sacrifice at 5 weeks of age.

### Reconstitution of germ-free mice

Fecal microbiota transfers (FMT) were performed by taking SPF feces from sex-matched adult WT mice and diluting 160 fold in sterile reduced PBS. Adult (6-10 weeks old) germ free (GF) mice were gavaged once with 150ul of solution.

### Cell lines

The mouse TLR4 reporter HEK293 cells (HEK-Blue mTLR4 cells) were engineered from the human embryonic kidney HEK293 cell line. Cells were used below 10 passages at 70-80% confluency and maintained at 37C with 5% CO_2_.

#### Colon cell isolation

To prepare single cell suspensions from colon tissue, mesenteric fat was removed, the tissue was cut longitudinally and washed in cold PBS to remove intestinal contents. The tissue was then placed in 15mL of pre-warmed HBSS+EDTA buffer (Hank’s Balanced Salt Solution [HBSS] supplemented with 5mM EDTA, 10% Fetal Bovine Serum [FBS] and 15mM HEPES) and incubated at 37°C, shaking at 250 rpm for 20 minutes. After incubation, the tubes were vortexed at the highest speed for 5 seconds and the tissue was removed and blotted on an absorbent wipe. This EDTA step was repeated. The tissue was then washed twice with 15mL of cold HBSS buffer (HBSS supplemented with 2% FBS and 15mM HEPES). Following the second wash, buffer was decanted, and excess liquid was removed with absorbent wipes. The tissue was digested in 5mL of digestion buffer (RPMI 1640 supplemented with 10% FBS, 15 mM HEPES, 100U/mL of DNase and 400U/mL of Collagenase VIII) for 35 minutes at 37°C, with shaking at 250 rpm. The digestion was stopped by adding 35 mL of cold R10 buffer. The tissue was passed and crushed through a 100μm filter and centrifuged at 2500 rpm for 10 minutes at 4°C. The cells were then re-suspended in R10 buffer and counted.

#### Flow cytometry

To assess cytokine production, colon LP cells were cultured in 24 well plates (1.5x 10^6^ cells/well) in T cell media (RPMI+ β-Mercaptoethanol (1.272ul/ml), Penicillin-Streptomycin (10ul/ml), and 10% FBS). To assess in vivo IL-17 and IL-1β production, cells were incubated with Golgi stop (0.67ul/ml) alone for 4 hours. To stimulate ex vivo production of IL-17, colon cells were stimulated with IL-1β (10ng/ml), IL-23 (10ng/ml) and Golgi Stop for 4 hours in T cell media. For IL-10 detection, colon cells were stimulated with PMA(50ng/ml), ionomycin (1ug/ml) and Golgi stop (0.67ul/ml). For stimulating ex vivo production of IL-1β, colon cells were stimulated with ATP (3mM) and LPS (100ng/ml). To test role of IL-10, IL-10 (10ng/ml) was added at the start of the culture. For in vitro stimulation with heat-killed bacteria, colonic LP cells were purified first using a Percoll gradient and stimulated for 24 hours with 1:10 (bacteria to cells ratio) HK *C.difficile* or *S.aureus*. PP2 (5μM) or anti-IL-1R (20ng/ml) was added to the 24-hour cultures as indicated. After ex vivo isolation and culturing, cells were washed in PBS prior to staining. For extracellular staining, single cell suspensions were incubated with fixable viability dye in 100μL PBS (eBioscience), washed and incubated with anti-Fc receptor (clone 2.4G2, BD biosciences) for 10 minutes before adding fluorochrome-labeled antibodies at pre-determined concentrations in 100μL FACs buffer (PBS containing 2% FBS and 10mM HEPES) for 30 minutes on ice. For intracellular staining, cells were fixed after extracellular staining with the Intracellular fixation/ permeabilization buffer (eBioscience) according to the manufacturer’s instructions followed by antibody labeling in 100μL permeabilization buffer. After intracellular labeling, cells were washed and resuspended in 300μL FACS buffer. Data were acquired on an LSR Fortessa (BD Biosciences) and analyzed with FlowJo software.

#### Single-Cell Sequencing library prep and analysis of mouse colon

Colon LP cells were isolated as previously described. Single cell suspensions were first incubated with fixable viability dye in 100μL PBS (eBioscience) for 20 minutes and washed with PBS. Cells were next incubated with anti-Fc receptor (clone 2.4G2) in PBS for 20 minutes, and washed. Cells were stained with fluorochrome-labeled CD45 antibody (clone 104) in 200 μL FACs buffer for 30 minutes for sorting CD45+ cells from the LP. Cells were then incubated with Total Seq hashtag antibodies (Biolegend cat #155831, #155837,or #155843) with antibodies against mouse CD45 (clone 30-F11) and MHC I complex (clone M1/42) for multiplexing and Total Seq TCR γ/δ antibody (Biolegend cat #118145) for 30 minutes. Stained single cell suspensions were then sorted individually for viability dye negative and CD45 positive population. Cell clumps were removed by passing cell suspension through 40µm FlowMi strainer (Bel-Art, USA). Samples from the same groups were combined in equal proportions. Single cell capture and library preparation was performed with Chromium Next GEM Single Cell 3ʹ Reagents Kit v3.1 (10X Genomics, USA). Cell suspensions were loaded on a Chromium Next GEM Chip G (10X Genomics, USA) together with gel bead from Chromium Next GEM Single Cell 3ʹ GEM Kit v3.1 and captured on Chromium Controller (10X Genomics, USA) with recovery target of 1×104 cells. cDNAs were generated following the 10X Genomics protocol CG000315 their quality was checked with Bioanalyzer High Sensitivity DNA Kit (Agilent, USA). One quarter of the total cDNA was used to generate sequencing libraries using Library Construction Kit (10X Genomics, USA) and barcoded using Dual Index plate TT set A (10X Genomics, USA). Obtained libraries were double sided size selected using SPRI select beads (Beckman Coulter, USA) to enrich for fragments 300-800 base pairs long, centered at 450bp. Libraries were checked for quality with Bioanalyzer High Sensitivity DNA Kit and paired-end sequenced on Illumina NovaSeq 6000 S4 flowcells aiming to obtain 50,000 reads per cell. Using Chromium Controller, Chromium Next GEM chip G and Chromium Next GEM Single Cell 3ʹ Reagents Kit v3.1 (10X Genomics, USA) cell emulsion, with a target capture of 10,000 cells, were obtained for scRNA-seq and Cell Surface Protein library preparation. Following the 10X Genomics protocol CG000317_RevD, cDNA and DNA from cell surface protein Feature Barcode were amplified using Feature cDNA Primers 2 from 3’ Feature Barcode kit (10X Genomics, USA). After amplification step, samples were size-selected with SPRIselect beads (Beckman Coulter, USA), where cDNA bound to beads, while DNA from cell surface protein Feature Barcode remained in the solution. After magnetic separation, cDNA was eluted from beads and used for scRNA-seq libraries as described above, whereas supernatants were used for Cell Surface Protein Library construction. The cell surface protein Feature Barcode DNA was purified by additional round of SPRIselect beads precipitation and amplified with primers from the Dual Index plate NT set A (10X Genomics, USA). Library quality was assessed with Bioanalyzer High Sensitivity DNA Kit and paired-end sequenced on Illumina NovaSeq 6000 S2 flowcells aiming to obtain 40,000 reads per cell.

For preprocessing, samples were first demultiplexed with Cell Ranger^65^ (v7.1.0) using the mm10-2020-A reference transcriptome. Downstream analyses were performed with the Seurat R^66^ package (v5.1.0), including cell demultiplexing with the HTODemux^67^ function. ADT counts of protein TCR antibody tag were included in the analysis. Using Seurat, Filtered count matrices were log-normalized, highly variable features were identified, and data were scaled. Separate experiments were integrated using reciprocal PCA. For two-dimensional data visualization, we performed UMAP based on the first 30 dimensions of the rpca data reduction. Cells were clustered using graph-based clustering as implemented in Seurat, based on the first 20 rpca dimensions with a resolution of 0.5. Cells with more than 8% mitochondrial reads were excluded from the dataset. Ambient RNA contamination was corrected using SoupX^68^ in all samples. Putative doublets were identified and removed by combining two criteria: cells with abnormally high UMI counts or number of detected features, and cells co-expressing markers from distinct immune cell populations. This marker-based approach was applied following initial clustering and cell type annotation. Differential neighborhood abundance (DA) analysis between different conditions was performed using the miloR^69^ package (Bioconductor). A k-nearest neighbor (kNN) graph was constructed in PCA space using the first 30 principal components with k = 20 neighbors per cell. Cell neighborhoods were defined by sampling 10% of cells as neighborhood seeds. P-values were corrected for multiple testing using Milo’s graph-aware spatial false discovery rate (SpatialFDR). Differential gene expression analysis between the conditions was performed using DESeq2^70^. Significantly differentially expressed genes (DEGs) (p-value < 0.05) and were identified and visualized using heatmaps.

#### Single-cell RNA-seq library construction and data processing for mouse thymus

Single-cell suspensions from thymi of eight postnatal day 0 mice were labeled with TotalSeq-C 5′ barcoded hashing antibodies (BioLegend) to enable sample demultiplexing, according to the manufacturer’s instructions, prior to FACS sorting. Libraries for sc RNA sequencing were generated from 1*104 FACS-pool sorted γδ T cells using the Chromium Single Cell 5’ Library Gel Bead and Construction kit (10x Genomics, CA, USA) according to the user guidelines (v2 Chemistry. Agilent Bioanalyzer High Sensitivity DNA chips were used to check quality control read-outs of sc RNA-seq libraries using a Bioanalyzer 2100 machine (Agilent Technologies). Indexed libraries were pooled and sequenced on the Illumina NovaSeq 6000 from the BRIGHTcore (Brussels Interuniversity Genomics High Throughput core) platform. Single-cell RNA sequencing was performed using 10x Genomics with 5’ feature barcoding technology to multiplex cell samples from different subjects. CellRanger (v7.1.0) software from 10x Genomics was used to demultiplex and map sequencing reads against reference. Count matrices were loaded into R using‘read10x’ function from Seurat R package. All downstream analyses were conducted using Seurat v5.0.0 in R v4.3.2. Low-quality and doublet cells were filtered out using the minimum cutoffs nFeature_RNA > = 200, % mitochondrial gene >5%. ScaleData function was used to normalize and stabilize variance in the datasets regressing out percent mitochondria genes, percent ribosomal genes, cell cycle scores, nFeatureRNA and TRDV TRGV genes. Principal component analysis (PCA) was subsequently performed using the RunPCA function with 30 princi”al c’mponents. To account for batch effects in integration of datasets, the Harmo”y al’orithm was applied using the RunHarmony function. The Harmony embeddings were extracted using the Embeddings function. Dimensionality reduction was then carried out using the RunUMAP function, based on the top 10-20 Harmony components to generate a two-dimensional UMAP projection for visualization. The nearest-neighbor graph was constructed with the FindNeighbors function, and clustering was performed using the Louvain algorithm via the FindClusters function at a resolution of 0.5. Visualization of cell clusters and feature expressions was carried out using DimPlot and FeaturePlot, respectively. Differential gene expression analysis was performed using the FindAllMarkers function to identify marker genes for each cluster, filtering for genes with a minimum expression threshold of 0.25 and log fold change ≥ 0.25. Single-cell gene signature enrichment scores for IL17-commitment and IFNγ-commitment were calculated using the ‘AddModuleScore’ function of Seurat using a list of genes previously described^38^.

#### Single cell sequencing of human fetal thymus

Single-cell RNA sequencing data of human fetal γδ thymocytes were obtained from the Gene Expression Omnibus (GEO) under accession number GSE180059^38^. IL1R1 expression in γδ thymocytes was visualized using the FeaturePlot function from the Seurat package.

#### Neonatal γδ T cell adoptive transfer

Thymi were collected from γδTCR+/+ mice at post-natal day 1. Thymi were crushed and cells were washed in R10 (RPMI+ 10% FBS+ 1%P/S + 1% glutamine + 15mM HEPES), counted and resuspended in PBS at 20 x 10^6^ cells/ml. 1-2 day-old recipients were first removed from their breeder cage and placed on ice for 1-2 minutes to “anesthetize” mice. 50ul of the solution was injected through the temporal vein. Mice were then harvested 3 weeks and 8 weeks post-injection to quantify the γδ17 T cell response.

#### Isolation and growth of unknown C. difficile strain from McGill microbiota

Fresh feces were diluted in PBS and plated onto selective *C. difficile* Brucella Agar (CDBA) plates in anaerobic conditions. CDBA plates were prepared by dissolving Brucella broth powder (28g/L), sodium bicarbonate (100mg/L), mannitol (6g/L), phenol Red (30mg/L), and agar (15g/L) in ultrapure water (Milli-Q), followed by autoclaving. After cooling, the media was supplemented with sodium taurocholic acid (0.5g/L), lysozyme (5mg/L), cycloserine (0.5g/L) and cefoxitin (16mg/L). Following incubation, a single colony of *C. difficile* was collected and flash frozen in liquid nitrogen. For strain identification, DNA was extracted using the Dneasy QIAGEN blood and tissue kit. The unknown *C. difficile* strain was sequenced using Nanopore. Contamination check was done using Kraken. The genome assembly was done using FLYE. Genome quality assessment was performed using Check M/QUAST. Strain was identified using FASTANI genome comparisons, toxin expression and MLST /typing.

#### Isolation of C. difficile spores and colonization

All culturing was performed at 37°C in a vinyl anaerobic chamber (Coy Laboratories, Grass Lake, MI, USA), maintained with a gas mixture of 3% H_2_, 10% CO_2_, 87% N_2_. A single colony of *C. difficile* was inoculated into brain heart infusion (BHI) broth (37 g/L) and grown for 48 hours. Subsequently, 200 µL of bacterial culture was spread onto a 70:30 sporulation agar plates using a sterile cell spreader. 70:30 sporulation plates were prepared by dissolving bacto-peptone (63 g/L), proteose peptone (3.5 g/L), ammonium sulfate (0.7 g/L), Tris base (1.06 g/L), BHI broth base (11.1 g/L), yeast extract (1.5 g/L), agar (15 g/L), and L-cysteine (0.5 g/L) in ultrapure water, followed by autoclaving. Plates were wrapped with parafilm and incubated for 21 days to allow for spore formation. For spore harvest, 2 mL of sterile ultrapure water was added to the surface of each plate, and growth was gently scraped using a sterile inoculating loop. The resulting suspension was centrifuged at 20 000 *g* for 10 min, and the pellet was resuspended in ultrapure water. Samples were stored at 4 °C for 3 days to facilitate spore release from vegetative cells. Lysis buffer (lysozyme, 30 mg/mL; Triton X-100, 10 µL/mL; in 10 mM Tris-HCl, pH 8.0) was added at 1:10 (v/v), and samples were incubated at 37°C for 2 hours, followed by sonication on ice. Spores were purified by density separation using a 50% (w/v) sucrose gradient and centrifugation at 5 000 *g* for 20 min. The supernatant was discarded, and the spore pellet was collected. Spores were resuspended in ultrapure water and washed five times by centrifugation at 14 000 *g* for 5 min. The final pellet was resuspended in ultrapure water and stored at 4 °C until use. Viable spores were enumerated by plating onto BHI+TCA plates and CFU quantification after 48h. BHI-TCA was prepared by autoclaving BHI broth base (37g/L), yeast extract (5g/L), agar (15g/L) in ultrapure water and supplementing with taurocholic acid (0.5g/L), and L-cysteine (0.5g/L) after cooling. Mouse colonization was detected by collecting feces at indicated times, diluting in PBS in anaerobic conditions and plating on CDBA plates. Colonies were counted following a 48-hour incubation period.

#### 16S sequencing of fecal microbiota

Fecal pellets were collected from CR and McGill mice and flash frozen in liquid nitrogen. DNA was extracted using QIAGEN PowerSoil kit, following manufacturer’s instructions. DNA was quantified using Nanodrop and sent for sequencing by Genome Quebec. 16S V3-V4 regions were amplified prior to sequencing using primers: 341F – CCTACGGGNGGCWGCAG and 805R – GACTACHVGGGTATCTAATCC. Each amplicon was sequenced and the sequencing data was processed and analyzed using the QIIME2 pipeline. DADA2, implemented as a QIIME2 plug-in, was used for sequence quality filtering^65^. For diversity metrics including the Faith phylogenetic diversity and UniFrac distances, a rooted phylogenetic tree was generated. Statistical analyses were carried out using R (version 3.6.0). Shannon index and richness were calculated using the package vegan. For differential abundance analysis, LeFSE was used with a significance threshold of 0.05. Plots were generated using the package ggplot2.

#### H&E Staining

Formalin-fixed paraffin-embedded blocks were cut to 5μm on a microtome, as described for immunofluorescence. After drying, slides were hydrated by 3×3 min washes in xylene, 3×3 min in ethanol and 1 min in 95% ethanol. Slides were placed in Mayer’s hematoxylin for 10 min followed by washing and blueing in an acidic bath. Eosin counterstain was done for 30 sec followed by 2×2 min washes in 90% ethanol and 3×2 min washes in xylene. Slides were mounted in a xylene-based mountant (Cytoseal; Thermofisher) and imaged on an Olympus BX50 microscope at 20X magnification.

#### Immunofluorescence

ZO-1 quantification was conducted with immunofluorescence on formalin-fixed paraffin-embedded tissues (FFPE). Freshly dissected tissues were cut longitudinally, washed in cold PBS, immediately prepared into a “Swiss roll” and placed in a histological cassette. Tissue was fixed in 10 % formalin for 24 h before being rinsed and transferred to 70 % ethanol. Sections were paraffin embedded by the histological core at the RI-MUHC and cut into 5 μm sections using a microtome (Leica). Sections were deparaffinized in sequential xylene washes followed by serial hydration steps in descending concentrations of ethanol. Antigen retrieval was done in citrate buffer for 20 min in a steamer followed by rest in the hot citrate buffer for 20 min at room temperature. Slides were washed 3 times for 5 min in PBS before blocking with 5% donkey serum with Fc block in PBS for 3 h. Samples were incubated with the primary antibody (indicated in key resources table) for overnight at room temperature followed by a secondary anti-rabbit IgG AF555 antibody stain for 2 h at room temperature in a hydration chamber. Slides were mounted in Prolong diamond antifade mountant (Invitrogen, Cat#P36970) and allowed to cure at room temperature overnight before imaging on Zeiss confocal microscope (LSM880). Images analysis was done in ImageJ (NIH) and in QuPath.67. ZO-1 quantification was conducted in ImageJ (NIH) as the number of positive ZO-1 pixels per complete villi. Each data point represents an average of at least 5 complete villi per mouse and represented as percentage positive ZO-1 area per total villi.

#### RNAScope

Slides were prepared as described for immunofluorescence. RNAScope was performed according to manufacturer instructions (ACD bio, Cat#322310) using the RNAscope 16S bacterial probe bacterial (Cat#467441). Images were acquired with an Axio Imager M2 microscope (Zeiss) at 20X magnification. Images were quantified using the pixel count function of Qupath^72^, and represented as the total positive pixel area along the colonic crypt-villus axis. Each data point represents the average pixel intensity from at least 3 independent colonic region per mouse.

#### Gene expression analysis

RNA was extracted from 0.5 cm of tissue using Tri reagent (Sigma-Aldrich) following manufacturer’s instructions, with an additional overnight ethanol precipitation step. Briefly, cleaned RNA was resuspended in 90 μl of water with 10 μl of 3M ammonium acetate and combined with 300 μl of ice-cold ethanol and left at-20°C for 18 h. Purified RNA was centrifuged, washed in 70% ethanol and resuspended in a final volume of RNase/DNase-free water. Reverse transcript was performed with AdvanTech 5x Reverse Transcription Mastermix (Cat# AD100-31404) and quantitative PCR was conducted using Advance Tech QPCR Mix (Advantech, Cat# AD100-21 402) and analyzed relative to HPRT internal controls using the 2-ΔΔCT method. All primer pairs are listed in the key resources table.

#### Human Vγ9Vδ2 T cell stimulation

CBMC (cord blood mononuclear cells) were stimulated with 10 uM zoledronate (Zometa, Novartis) in the presence or absence of recombinant human IL-1β (at 10 ng/ml) ((R&D Systems) in 14 ml tubes at a final concentration of 2.10e6 cells/ml. Intracellular staining for IFN-γ and IL-17 was performed using the Cytofix/Cytoperm kit (BDBiosciences). For the detection of IFN-γ and IL-17, monensin (2 mM, Sigma-Aldrich, Bornem, Belgium) was added the last 4 hours of the PBMC and CBMC cultures. Cells were run on the CyAn flow cytometer equipped with 3 lasers (405 nm, 488 nm, 633 nm) and data were analyzed using Summit 4.3 (DakoCytomation).

## ACKNOWLEDGEMENTS

We thank the staff of the animal, histopathological, and immunophenotyping platforms at the Research Institute of the McGill University Health Centre. Special thanks to Cynthia Faubert, Catherine Hudon and Caroline Monat for their assistance with the germ-free animal studies and anaerobic bacterial cultures. Thanks to the McGill Interdisciplinary Initiative in Infection and Immunity for financial support to establish the Gnotobiotic Animal Research Platform at the RI-MUHC. We also appreciate the Meakins-Christie Laboratories administration and Severine Audusseau for technical support. B.T. is the recipient of a Canadian Institutes of Health Research (CIHR) Doctoral Fellowship and Fonds de Recherche du Quebec Doctoral award. S.W. is the recipient of a Canadian Institutes of Health Research (CIHR) Postdoctoral Fellowship. I.L.K. is a recipient of a Fonds de Recherche du Quebec Scholar Award. This work was supported by a CIHR Project Grant (PJT-183685) to I.L.K.

## AUTHOR CONTRIBUTIONS

Conceptualization, B.T. and I.L.K.; methodology, B.T., S.W., L.C., A.S., K.O., E.M., Y.T., P.L., G.F., Benoit C., S.G., D.V., Bastien C., A-E.S., and I.L.K.; Software, B.T., L.C., A.S., Y.T.; formal analysis, B.T., S.W., L.C., L.K., A.S., K.O., E.M., Y.T., and I.L.K.; Investigation, B.T., S.W., L.K., K.O., E.M., Y.T., G.F., and I.L.K.; Resources, A.S., B.J.S., P.L., Benoit C., S.G., D.V., Bastien C., A-E.S., and I.L.K.; Writing original draft, B.T., and I.L.K.; Writing – review & editing, all authors; Visualization, B.T., S.W., L.C., and I.L.K.; funding acquisition, I.L.K.

---

**Figure S1.** Immune activation in the early life colon is mediated by. γδ**17 T cells. (A)** Number of IL-22+ γδ T cells, Th17 and ILC3s in the colonic lamina propria at indicated ages. **(B)** Percent of IL-17+ γδ T cells in the proximal, mid and distal colon at weaning. **(C)** Percent of IL-22+ of colonic LP γδ T cells of mice at indicated ages following 4 hour stimulation with 10ng/ml IL-1β and 10ng/ml IL-23. **(D)** ELISA results for quantification of IFNγ, IL-13 and IL-17 in total colon lysates of weanling and adult mice. **(E)** Percent of IFNγ+ CD4+ cells in the colon, small intestine and lungs of weanling and adult mice. **(F)** UMAP of TCRδ-specific antibody tagging in total colonic LP CD45+ cells. **(G-H)** Differentially expressed genes (G) and GO analysis pathways (H) in γδ T cells of weanling and adult mice. (A, B, C, D, E) Each dot represents cells from an individual mouse. Data are represented as mean with SD. Graphs are pooled data from at least two independent experiments with atleast n = 3 in each experiment. (B, D, E) Data were analyzed using a two-way ANOVA (∗p < 0.05, ∗∗p < 0.01, ∗∗∗p < 0.001, ∗∗∗∗p <0.0001).

**Figure S2.** Immunophenotyping and scRNA sequencing analysis of colonic CD45+ cells from weanlings and adults. (A, **B)** Total number of neutrophils and monocytes in the colon LP of 3-week-old littermate TCRδ+/-and TCRδ-/-mice. **(C, D)** Total number of neutrophils and monocytes in the colon lamina propria of control mice and mice treated with anti-IL-17 in vivo antibody from 2-3 weeks of age. **(E, F)** Top 30 expressed genes in the monocytes (E) and the neutrophils (F) from 3-week-old colon, after removal of ribosomal and mitochondrial genes. **(G)** Normalized expression of *Il1b* transcripts in different immune population in total CD45+ cells of the colon LP. **(H, I)** Genes upregulated in different Treg subsets (H) and corresponding UMAP (I) in total immune cells of colonic LP. **(J)** UMAP indicating *Rorc* expression in weanling and adult Tregs (A-D) Each dot represents cells from an individual mouse. Data are represented as mean with SD. Graphs are pooled data from at least two independent experiments with atleast n = 3 in each experiment. Mann-Whitney non-parametric unpaired T test (∗p < 0.05, ∗∗p < 0.01, ∗∗∗p < 0.001, ∗∗∗∗p <0.0001).

**Figure S3.** γδ**17 T cell activation and IL-1**β **production is microbiota-dependent. (A-B)** Number of γδ T cells (A) and percent of Ki67+ γδ T cells (B) in weanling SPF and GF mice. **(C)** Percent of IL-22+ γδ T cells in the colon LP following GF reconstitution at indicated times, n = 4. **(D)** Percent of IL-22+ γδ T cells in the colon LP 1 week following GF reconstitution with CR and McGill microbiota. **(E)** Percent of IL-22+ γδ T cells in the colon LP of 3-week-old pups with CR or McGill SPF microbiotas. **(F, G)** UMAP (F) and MiloR differential abundance analysis (G) of single cell sequencing of CD45+ immune cells in CR and McGill weanling mice. (H, I) Percent of pro-IL-1β+ myeloid cells (H) and percent of neutrophils (I) in the colon of mice with CR microbiota or CR + McGill microbiota at indicated ages. (A-E, H, I) Each dot represents cells from an individual mouse. Data are represented as mean with SD. Graphs are pooled data from at least two independent experiments with at least n = 3 in each experiment unless otherwise indicated. (C-E) Data were analyzed using Kruskal-Wallis ANOVA test or (A, B) Mann-Whitney non-parametric unpaired T test or (H, I) using 2-way ANOVA test (∗p < 0.05, ∗∗p < 0.01, ∗∗∗p < 0.001, ∗∗∗∗p <0.0001).

**Figure S4.** McGill microbiota-associated *C. difficile* is related to an outbreak strain. (A, **B)** FastANI comparison of the unknown strain of *C. difficile* to ATCC strain VPI 10463 and other known strains organized by similarity (A) and clades (B). **(C)** Protein level of IL-1β following 24-hour in vitro stimulation of gut LP immune cells with HK *C. difficile,* n = 3. **(D, E)** Number of colonic LP γδ T cells (D), and percent of IL-22+ γδ T cells (E) in the colon of control CR pups, CR pups colonized with *C. difficile* spores and their littermates. **(F-I)** Percent of Ki67+ γδ T cells (F), percent of IL-22+ γδ T cells (G), percent of neutrophils of CD45+ cells (H) and percent of IL-1β+ myeloid cells (I) in the colon of CR control pups and pups of *C. difficile* colonized breeders. **(J, K)** Percent of Rorγt+ γδ T cells in the epithelial fraction of mice at indicated ages (J) and comparing SPF and GF mice (K). (C-K) Each dot represents cells from an individual mouse. Data are represented as mean with SD. Graphs are pooled data from at least two independent experiments with atleast n = 3 in each experiment, unless indicated. (D, E, J) Data were analyzed using Kruskal-Wallis ANOVA test or (C, F, G, H, I, K) Mann-Whitney non-parametric unpaired T test (∗p < 0.05, ∗∗p < 0.01, ∗∗∗p < 0.001, ∗∗∗∗p <0.0001).

