## Supplementary figures and images for "Early life γδ T cell activation enforces intestinal barrier integrity during intergenerational *C. difficile* colonization"

### Supp Figure 1

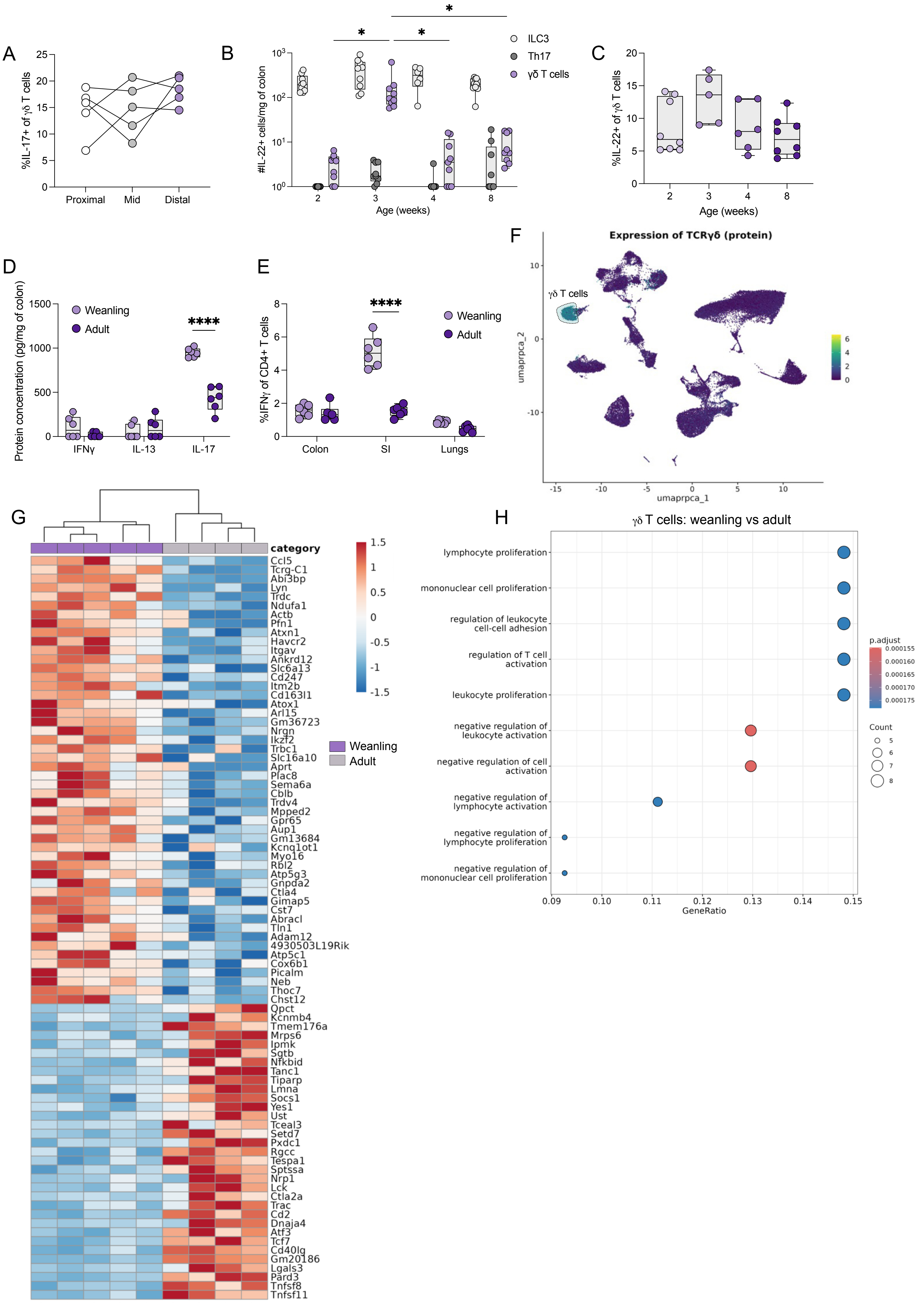

### Supp Figure 2

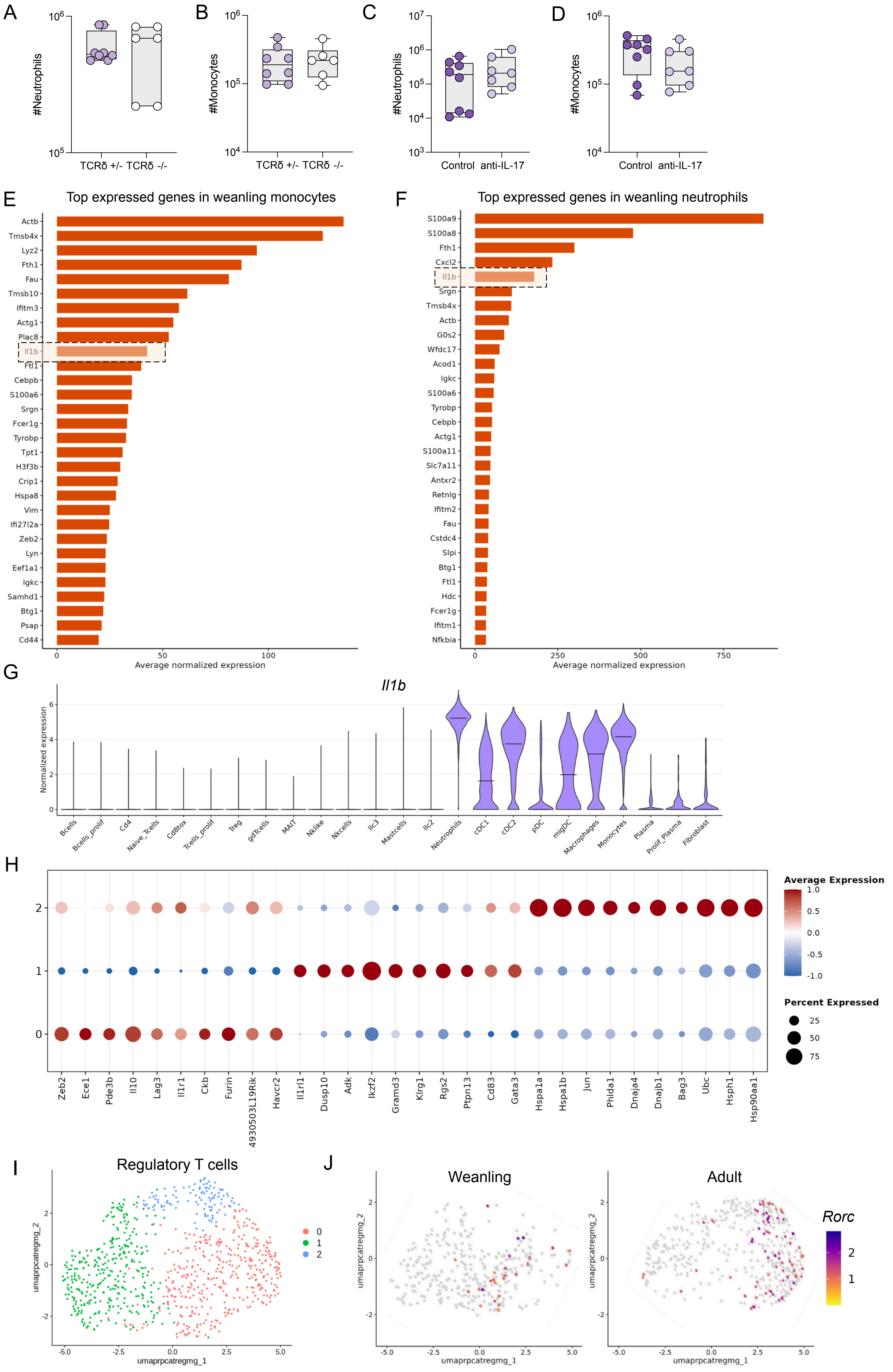

### Supp Figure 3

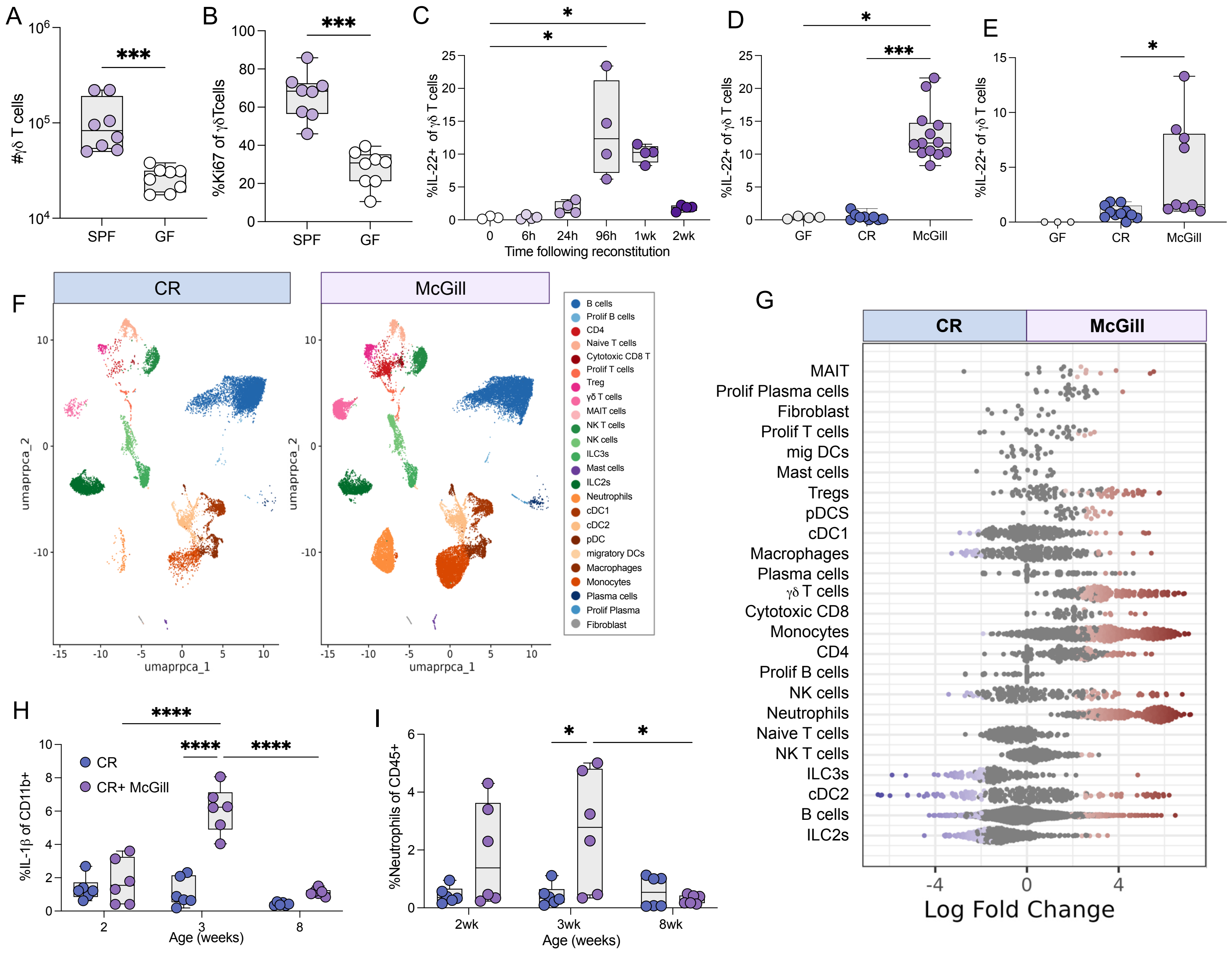

### Supp Figure 4

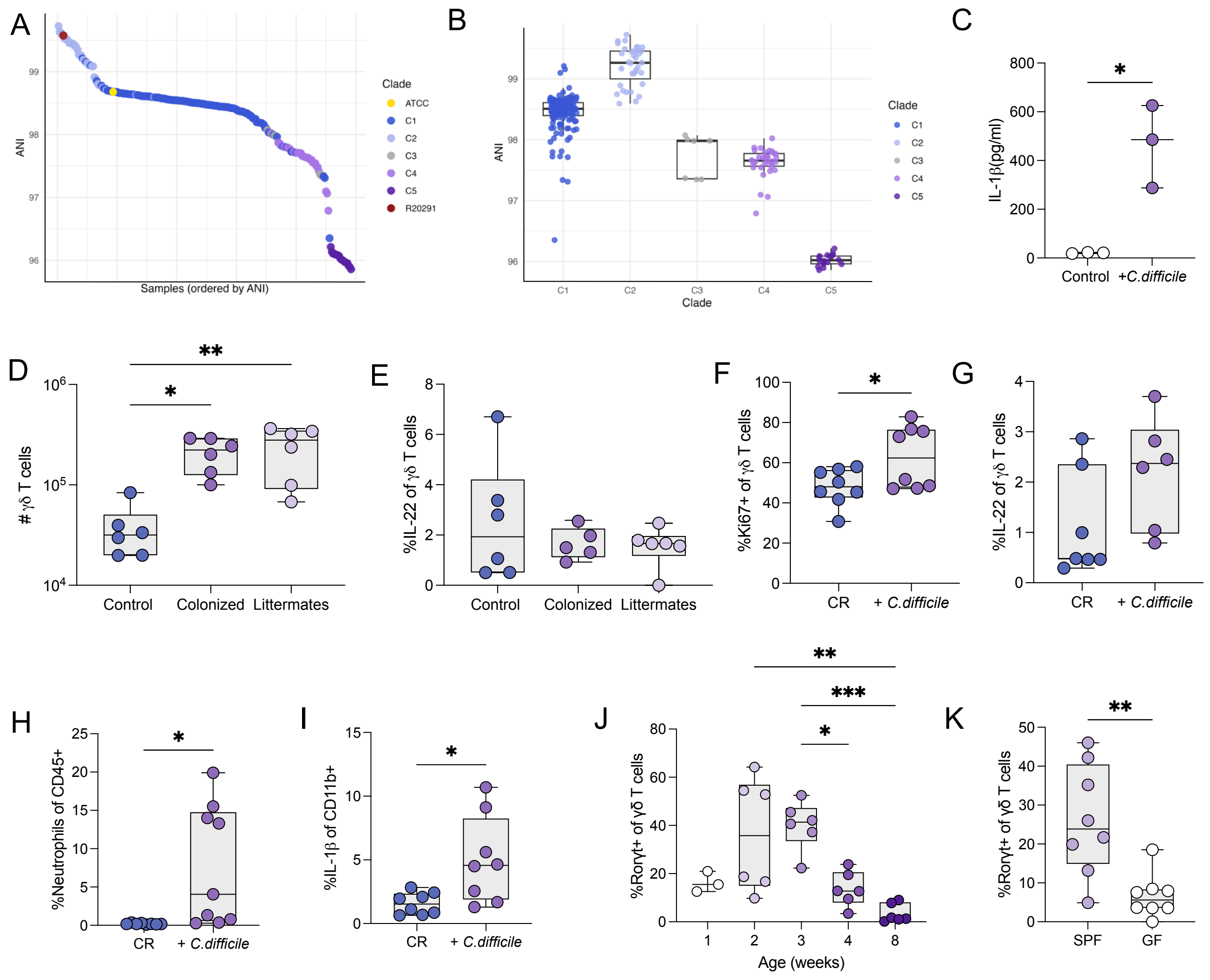
